# Fatty acyl-CoA reductase FAR1 is essential for testicular seminolipid synthesis, spermatogenesis, and male fertility

**DOI:** 10.1101/2024.12.26.630343

**Authors:** Ayano Tamazawa, Tatsuro Naganuma, Kento Otsuka, Tenga Takahashi, Takayuki Sassa, Akio Kihara

## Abstract

Seminolipids are testis-specific ether glycolipids that are important for spermatogenesis. The fatty alcohol (ether-linked alkyl moiety) in ether lipids is generated from an acyl-CoA by fatty acyl-CoA reductase (FAR). To date, the diversity of the alkyl and acyl moieties in seminolipids, the specific stage of spermatogenesis during which seminolipids are produced, and the FAR isozyme (FAR1 or FAR2) involved in the synthesis of the alkyl moieties have remained largely unclear. Here, we demonstrated that *Far1* is expressed in the mouse testis via quantitative RT-PCR analysis, whereas *Far2* was barely detectable. *In situ* hybridization and quantitative RT-PCR analysis of spermatogenic cells separated via FACS revealed that *Far1* is expressed in spermatogonia, spermatocytes, and spermatids. We generated *Far1* knockout (KO) mice and found that male *Far1* KO mice were infertile. In these mice, sperms were absent in the epididymides and the testes were small, with multinucleated cells and vacuoles in the seminiferous tubules. LC-MS/MS analysis showed that the vast majority of seminolipids (> 90%) in wild-type mouse testes contained C16:0 in both the alkyl and the acyl moieties. Seminolipids were present in all subclasses of spermatogenic cells in wild-type mice, but they were absent in *Far1* KO mice. Instead, the production of non-ether, diacyl-type sulfogalactosyl lipids (sulfogalactosyl diacylglycerols) was induced in *Far1* KO mice. In conclusion, the alkyl and acyl moieties of seminolipids in the testis are low in diversity, and *Far1* is essential for seminolipid synthesis and spermatogenesis.

## Introduction

Sperms have unique properties not found in other cells, including a haploid genome, the ability to move with the aid of a flagellum, and the capacity to fertilize eggs. To acquire these properties, sperms are produced via complex, spatiotemporally regulated processes involving the differentiation and maturation of spermatogenic cells. The production and functioning of spermatogenic cell- and sperm-specific proteins and lipids are essential for these processes. The testis is composed of seminiferous tubules, which produce sperms, and interstitial connective tissue containing Leydig cells, which produce sex hormones (1). Sperms produced in the seminiferous tubules are transported via the efferent ductule of the testis to the epididymis, where they become highly motile and acquire the ability to fertilize. The epithelium in the seminiferous tubule contains spermatogenic cells and Sertoli cells, which form a blood–testis barrier and support spermatogenesis physically and nutritionally (1, 2). Spermatogenic cells are further subdivided into spermatogonia, spermatocytes, and spermatids. The spermatogonium, located on the basal lamina of the seminiferous epithelium, is a stem cell that undergoes somatic division to self-replicate or differentiate into spermatocytes. Spermatocytes differentiate into haploid spermatids via meiotic division. The morphology of the spermatid is initially round, but it elongates as the acrosome and flagellum are formed. During the division of spermatogenic cells, cytokinesis is incomplete, resulting in the cells’ cytoplasm being connected by intercellular bridges (3). At the stage when the elongated spermatids release residual bodies, the spermatids separate and are released as sperms into the lumen of the seminiferous tubules. The residual bodies released are subsequently phagocytosed by Sertoli cells.

Seminolipids are lipids that exist specifically in spermatogenic cells (4). In mammals, most of the glycolipids are sphingolipids (glycosphingolipids), but seminolipids belong to the glycerolipids. A seminolipid has an alkyl moiety, an acyl moiety, and a sulfogalactose moiety at the *sn-*1, *sn-*2, and *sn-*3 positions, respectively (Fig. 1) (5). The synthesis of alkyl moieties in ether lipids, including seminolipids, occurs in the peroxisomes (6, 7). The fatty alcohol constituting the alkyl moiety of a seminolipid is produced by fatty acyl-CoA reductase (FAR), which reduces a fatty acyl-CoA to a fatty alcohol. In the final two steps of seminolipid synthesis, galactose is added to 1-alkyl-2-acyl-glycerol by ceramide galactosyltransferase (CGT), generating 1-alkyl-2-acyl-3-galactosyl-glycerol (GalAAG). Subsequently, sulfate is introduced into the galactose group of GalAAG by cerebroside sulfotransferase (CST). Both *Cgt* knockout (KO) mice and *Cst* KO mice lack seminolipids in their testes and exhibit defects in spermatogenesis (8, 9), demonstrating that seminolipids are indispensable for spermatogenesis.

**Figure 1.**
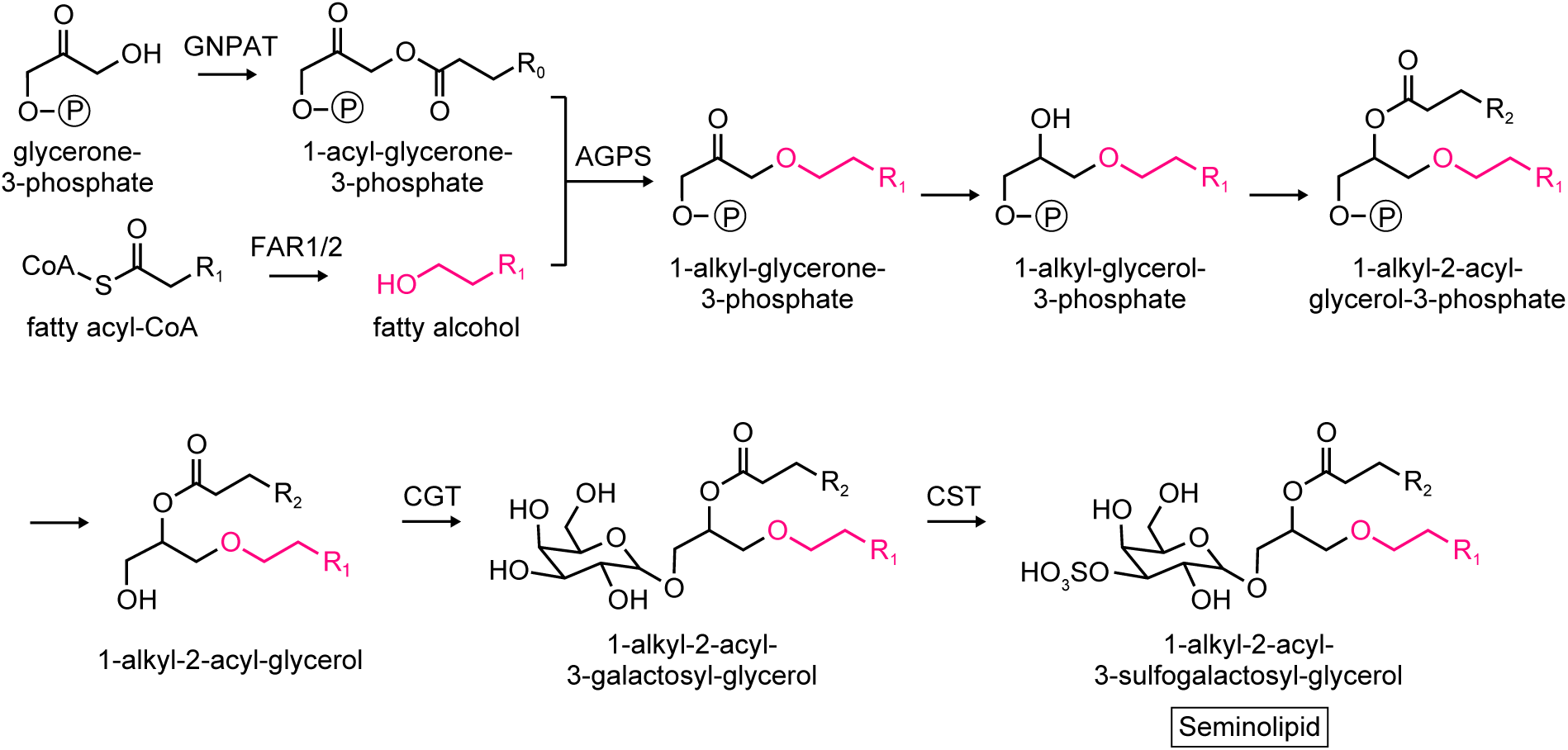
Synthetic pathway of seminolipids. In the first reaction, glycerone (dihydroxyacetone)-phosphate *O*-acyltransferase (GNPAT) esterifies a fatty acid to the *sn*-1 position of glycerone-3-phosphate. This fatty acid is then replaced by a fatty alcohol, which is linked to the *sn*-1 position via an ether bond, to form 1-alkyl-glycerone-3-phosphate. This reaction is catalyzed by alkylglycerone phosphate synthase (AGPS) and the fatty alcohol is generated from fatty acyl-CoA by the fatty acyl-CoA reductase (FAR). There are two FAR isozymes, FAR1 and FAR2, in mammals. 1-Alkyl-glycerone-3-phosphate undergoes reduction of a carbonyl group to a hydroxy group, esterification of a fatty acid to the *sn*-2 position, and removal of a phosphate group to form 1-alkyl-2-acyl-glycerol. Finally, a seminolipid is formed by the attachment of a galactose to 1-alkyl-2-acyl-glycerol by ceramide galactosyltransferase (CGT) and subsequent sulfation of the galactose moiety by cerebroside sulfotransferase (CST).

In mammals, there are two FAR isozymes: FAR1 and FAR2 (10). In mice, *Far1* is expressed in various tissues and organs, with the highest levels in the preputial glands, followed by the kidney, testis, and brain (10). In contrast, *Far2* is expressed in a relatively tissue- or organ-specific manner; its expression levels are highest in the eyelids (meibomian glands), followed by the skin, small intestine, and brain (10). FAR1 exhibits activity toward C16–18 acyl-CoAs, whereas FAR2 is active toward acyl-CoAs with a wider range of chain lengths (10, 11). Although it has been reported that the seminolipid species with C16:0 in both the alkyl and the acyl moieties (*O*-C16:0/C16:0, where *O-* indicates ether-linked) is abundant (12–14), the detailed composition of seminolipids in the testis remains undetermined. Furthermore, it remains unclear which FAR isozyme (FAR1 or FAR2) is involved in the synthesis of seminolipids and during which stages of spermatogenesis seminolipids are synthesized. Mutations in *FAR1* cause the inherited disease CSPSD (<u>c</u>ataracts, <u>s</u>pastic <u>p</u>araparesis, and <u>s</u>peech <u>d</u>elay) (15–18); however, spermatogenesis in patients with CSPSD has not been reported. During the preparation for this study, *Far1* KO mice were generated and reported to exhibit a severe reduction in plasmalogen levels in the testis and, similarly to *Cgt* KO and *Cst* KO mice, impaired spermatogenesis (19). However, seminolipids were not measured in that study, so the relative contributions of *Far1* and *Far2* in the synthesis of seminolipids were not determined.

In this study, we examined the expression of *Far1* and *Far2*, the alkyl/acyl composition of seminolipids, and the abundance of seminolipids at different stages of spermatogenesis in the testes of wild-type (WT) mice. We also independently generated *Far1* KO mice and analyzed their fertility, testicular histology, and lipid profile to reveal the role of *Far1* in seminolipid synthesis and spermatogenesis.

## Results

### *Far1* is expressed in the testis

To examine the contribution of the FAR1 and FAR2 isozymes to the synthesis of seminolipids, we first analyzed the mRNA levels of *Far1* and *Far2* in the testes of WT mice via quantitative RT-PCR and found that only *Far1*, and not *Far2* mRNA, was expressed in the testes (Fig. 2*A*). We then examined the distribution of *Far1* mRNA in the testes via *in situ* hybridization. The control experiment, which had no probe, generated no signals, but hybridization with the *Far1* probe stained cells lining the basal lamina and extending to the luminal center of the seminiferous tubules (Fig. 2*B*). This staining pattern was similar to that observed for the testis-specific histone gene *H2bc1*, which is expressed in spermatogenic cells, indicating that *Far1* is expressed in spermatogenic cells. *Far1* staining was not observed in the interstitial connective tissue outside the seminiferous tubules, where the Leydig cells exist.

**Figure 2.**
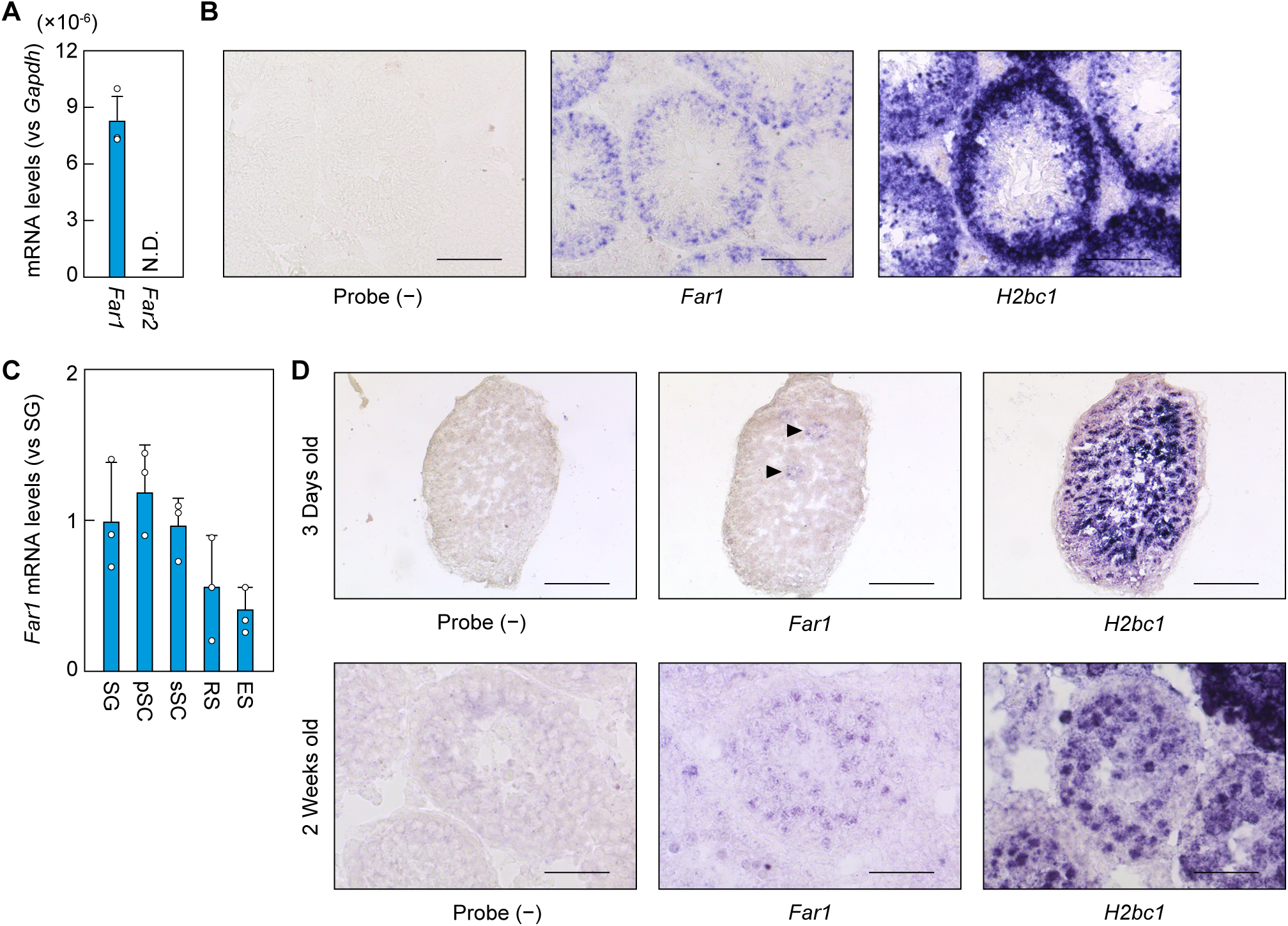
*Far1* is expressed in the testis. *A*, total RNAs were extracted from the testes of 8-week-old C57BL/6 mice and subjected to quantitative RT-PCR using specific primers for *Far1*, *Far2*, or the housekeeping gene *Gapdh*. Values presented are means + SD of each mRNA quantity relative to that of *Gapdh* (n = 3). N.D., not detected. *B*, fresh frozen sections were prepared from the testes of 6-month-old C57BL/6 mice and subjected to in situ hybridization using digoxigenin-labeled antisense RNA probes against *Far1* or *H2bc1*. Probe (−) represents a negative control experiment without probes. Scale bars, 100 µm. *C*, spermatogenic cells prepared from the testes of 8-week-old C57BL/6 mice were fractionated into spermatogonia (SG), primary spermatocytes (pSC), secondary spermatocytes (sSC), round spermatids (RS), and elongated spermatids (ES) by FACS. Total RNAs extracted from each fraction were subjected to quantitative RT-PCR using specific primers for *Far1* or the housekeeping gene *Gapdh*. Values presented are means + SD of *Far1* mRNA quantities relative to *Gapdh* (n = 3). *D*, fresh frozen sections were prepared from the testes of 3-day-old and 2-week-old C57BL/6 mice and subjected to *in situ* hybridization using digoxigenin-labeled antisense RNA probes against *Far1* or *H2bc1*. Probe (−) represents a negative control experiment without probes. Scale bars, 400 µm and 50 µm for 3-day-old and 2-week-old mice, respectively.

Next, to examine when *Far1* is expressed during the differentiation of spermatogenic cells, we isolated the spermatogonia, spermatocytes, round spermatids, and elongated spermatids from the testes of adult WT mice via FACS and measured the levels of *Far1* mRNA via quantitative RT-PCR. *Far1* was expressed in all spermatogenic cells (Fig. 2*C*). The mRNA levels in the spermatogonia and spermatocytes were approximately double those in the round and elongated spermatids.

During postnatal testis development, the spermatogenic cells in the seminiferous tubules at postnatal day three consist mostly of gonocytes, the precursors of spermatogonia, and include a small number of spermatogonia (20). By postnatal week two, spermatogonia and spermatocytes constitute about 80% and 20% of spermatogenic cells, respectively (21). Thus, cell types are limited at these stages and can be identified histologically. We examined *Far1* expression at these stages using *in situ* hybridization. At postnatal day three, *Far1* was expressed in only a small number of putative spermatogonia (Fig. 2*D*, arrowheads), and at postnatal week two, *Far1* was expressed in the spermatogonia and spermatocytes in the peripheral and central regions, respectively, in the seminiferous tubules (Fig. 2*D*). These results indicate that the expression of *Far1* starts when the spermatogonia are derived from gonocytes and continues during differentiation into spermatocytes and spermatids.

### 1-Alkyl and 2-acyl composition of seminolipids in the testis

Seminolipids in the testis consist predominantly of species with C16:0 as both their alkyl and their acyl moieties (*O*-C16:0/C16:0) (12–14). Smaller quantities of other species with C14:0– 18:0 alkyl or acyl moieties are also present (12, 13). To date, seminolipids other than the *O*-C16:0/C16:0 species have been analyzed via LC-MS, fast atom bombardment MS, and imaging MS (12–14). However, LC-MS and fast atom bombardment MS cannot distinguish between alkyl and acyl moieties in the *sn*-1 and *sn*-2 positions, respectively, and imaging MS has a poor quantitation capability. Thus, the precise alkyl/acyl composition of seminolipids has not yet been clarified. To address this issue, we analyzed seminolipids via LC-MS/MS, which can distinguish species with alkyl/acyl moieties of various chain lengths and degrees of unsaturation. In the analysis, the alkyl or acyl moiety was fixed at C16:0, whereas the other moiety was variable, with chain lengths of C14–26 and 0–6 double bonds (Fig. 3*A*). Consistent with previous reports (12–14), *O*-C16:0/C16:0 was the most abundant species, comprising 91% of the total seminolipids (Fig. 3*B*). The next most abundant species were *O*-C18:1/C16:0 (3%), *O*-C18:0/C16:0 (2%), and *O*-C16:0/C14:0 (2%). The following species were present in trace quantities (< 1%): *O*-C16:0/C15:0, *O*-C16:0/C16:1, *O*-C16:0/C17:0, *O*-C16:0/C18:1, *O*-C14:0/C16:0, *O*-C15:0/C16:0, and *O*-C16:1/C16:0.

**Figure 3.**
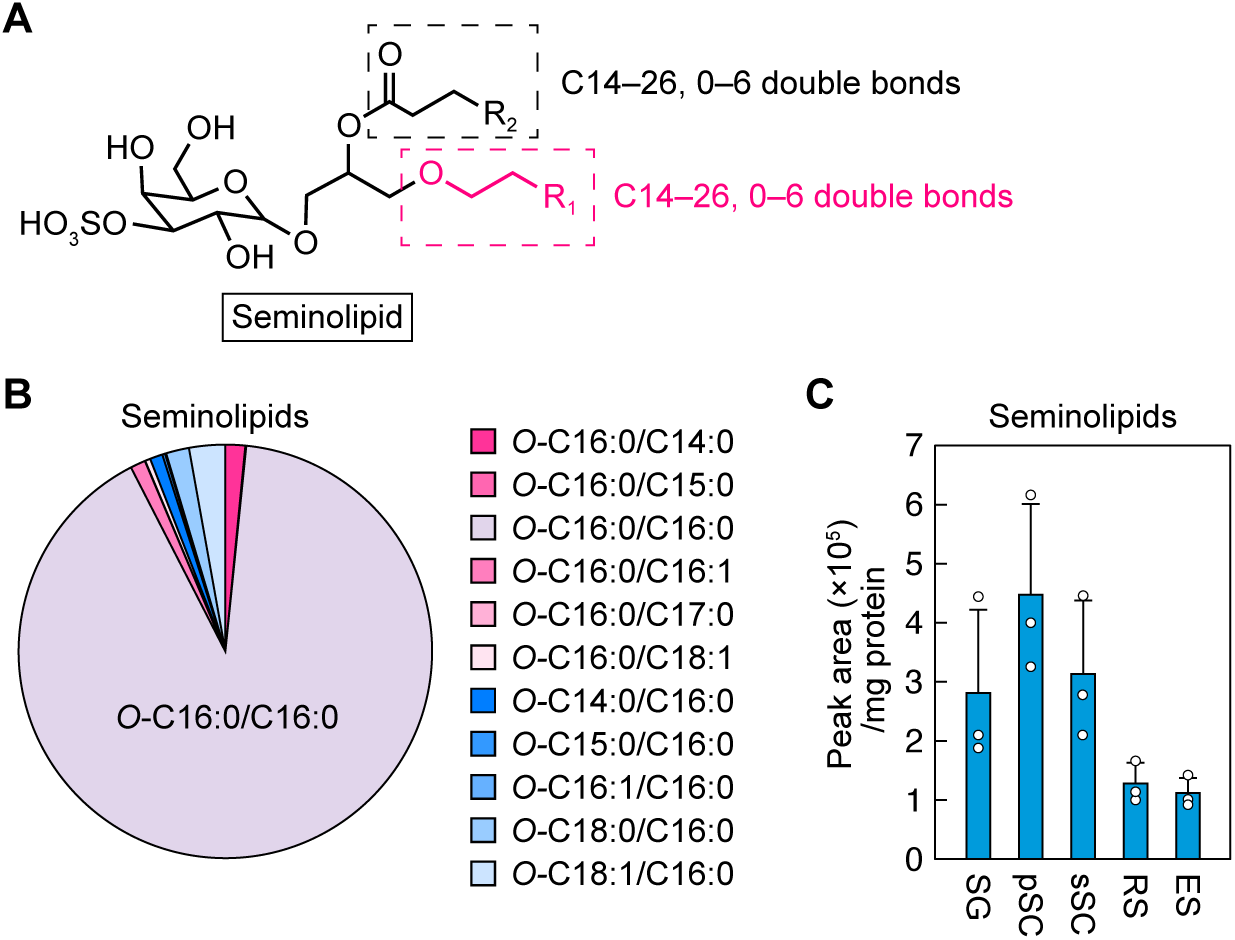
1-Alkyl and 2-acyl composition of seminolipids in the testis. *A*, the structure of a seminolipid. The ranges of chain lengths and degrees of unsaturation in 1-alkyl and 2-acyl moieties are noted, indicated respectively by black and red dashed rectangles. *B*, lipids were extracted from the testes of 8-week-old C57BL/6 mice and seminolipids were quantified via LC-MS/MS. The ratios of 11 species detected in the analysis are presented. *C*, spermatogenic cells prepared from the testes of 8-week-old C57BL/6 mice were fractionated into spermatogonia (SG), primary spermatocytes (pSC), secondary spermatocytes (sSC), round spermatids (RS), and elongated spermatids (ES) by FACS. Lipids were extracted from each fraction, and seminolipids were quantified via LC-MS/MS. Values presented are means + SD of total seminolipids in the indicated cell fractions (n = 3).

Next, to determine the stage of spermatogenesis at which seminolipids are present, testes from WT mice were dispersed and spermatogonia, primary and secondary spermatocytes, and round and elongated spermatids were separated via FACS. Lipids were extracted from each spermatogenic cell population and seminolipids were quantified via LC-MS/MS. The levels of seminolipids in spermatogonia and spermatocytes were similar and approximately double those in spermatids (Fig. 3*C*). The levels of seminolipids correlated well with *Far1* mRNA expression (Fig. 2*C*).

### Impaired spermatogenesis in *Far1* KO mice

We produced *Far1* KO mice using the CRISPR-Cas9 system to investigate the role of FAR1 in seminolipid synthesis and spermatogenesis. The guide RNA was designed to target the sequence downstream of the start codon in exon 3. The resulting *Far1* KO allele had a 22 bp deletion that encompassed the start codon (Fig. 4*A*). To obtain homozygous *Far1* KO mice, heterozygous *Far1* KO mice were interbred, and the offspring were genotyped at three weeks of age. Of 323 offspring, the numbers of WT, heterozygous KO, and homozygous KO mice were 96, 214, and 13, respectively; the proportion of homozygous KO mice (4.0%) was lower than the 25% expected under Mendel’s law (Table 1). To examine whether some homozygous *Far1* KO mice had died during embryonic development, we genotyped 122 embryos at embryonic day 18.5 (E18.5) and found that the numbers of WT, heterozygous KO, and homozygous KO mice were 37 (30%), 54 (44%), and 31 (25%), respectively (Table 2). This showed that the homozygous *Far1* KO mouse embryos were present at E18.5 in the proportion expected under Mendel’s law. We next examined the survival rate of WT and homozygous *Far1* KO mice (hereafter referred to as *Far1* KO mice) after birth. Of 12 newborn mice of each genotype, 11 WT (92%) and 6 *Far1* KO mice (50%) were alive on the day of birth, and nine WT (75%) and one KO mouse (8%) were alive on the following day (Fig. 4*B*). Thus, the *Far1* KO mice exhibited high early postnatal lethality, but not embryonic lethality. Production of more offspring resulted in some *Far1* KO mice that lived for up to four weeks after birth, with some of those even surviving more than one year.

**Figure 4.**
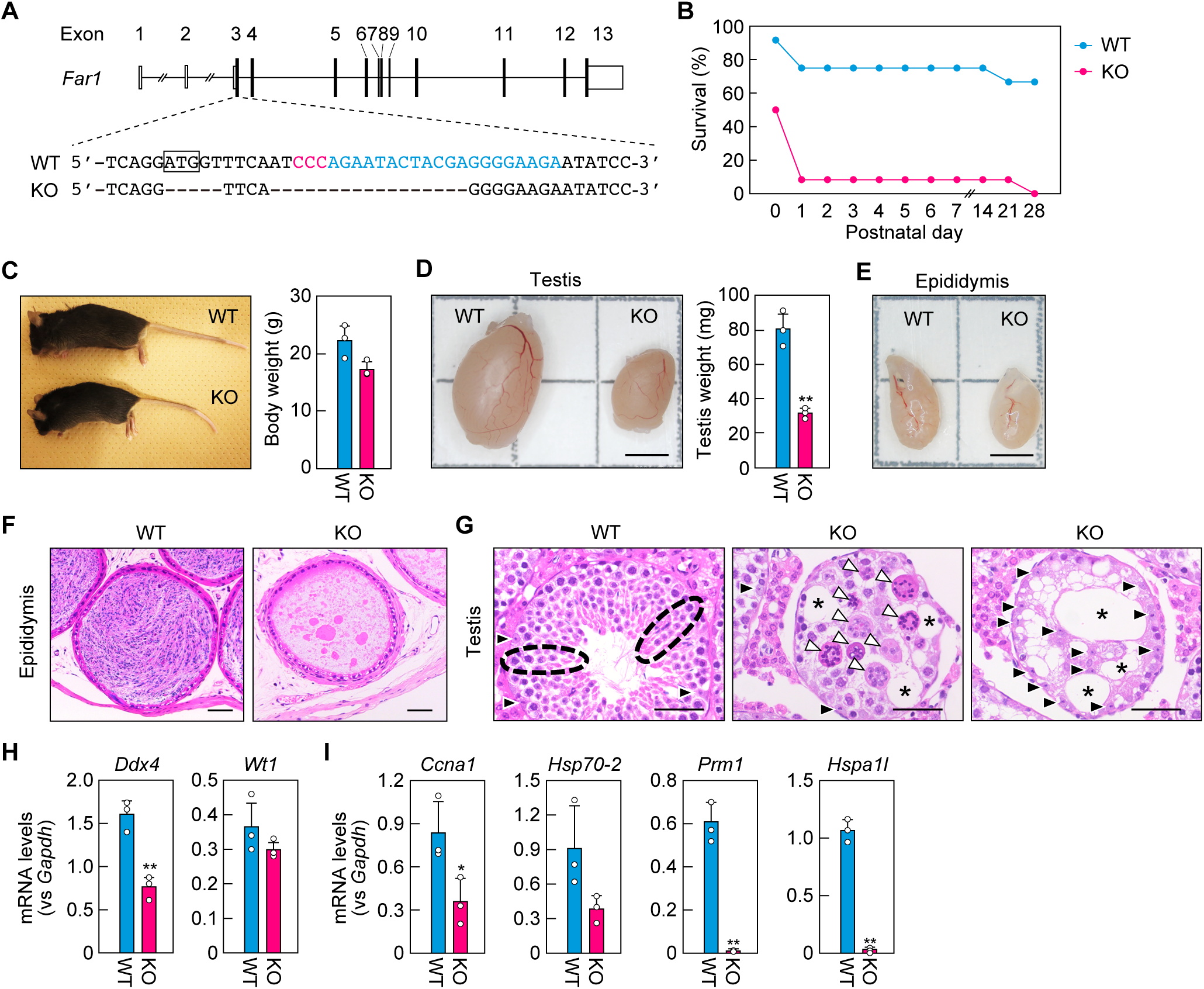
Impaired spermatogenesis in *Far1* KO mice. *A*, generation of *Far1* KO mice using the CRISPR-Cas9 system. The gene structure of *Far1* (coding regions and untranslated regions in black and white, respectively) is shown, along with the nucleotide sequences of WT and *Far1* KO mice around the guide RNA sequence (blue) and the protospacer-adjacent motif sequence (red). The box indicates a start codon. *B*, newborn WT and *Far1* KO mice (n = 12 for each genotype) were monitored at the indicated ages and the survival rates were calculated. *C*, the appearance and body weights of 8-week-old male WT and *Far1* KO mice. Values presented are means + SD (n = 3). *D*, the appearance and weights of the testes of 8-week-old male WT and *Far1* KO mice. Values presented are means + SD (n = 3). Statistically significant differences are indicated (Welch’s *t*-test; \*\**p* < 0.01). Scale bar, 3 mm. *E*, the appearance of epididymides of 8-week-old male WT and *Far1* KO mice. Scale bar, 3 mm. *F* and *G*, images of paraffin sections of the epididymides (*F*) and testes (*G*) from 7-month-old WT and *Far1* KO mice stained with hematoxylin and eosin. Scale bars, 50 µm. Black dashed ovals: examples of differentiating spermatogonia, spermatocytes, and spermatids, which were aligned from the basal lamina toward the center of the seminiferous tubules; black arrowheads: Sertoli cells; white arrowheads: multinucleated cells; asterisks: vacuoles. *H* and *I*, total RNAs extracted from the testes of 8-week-old male WT and *Far1* KO mice were subjected to quantitative RT-PCR using specific primers for *Ddx4* (pan-spermatogenic cell marker), *Wt1* (Sertoli cell marker), *Ccna*1 and *Hsp70-2* (primary spermatocyte markers), and *Prm1* and *Hspa1l* (spermatid markers), or the housekeeping gene *Gapdh*. Values presented are means + SD of mRNA quantities relative to *Gapdh* (n = 3). Statistically significant differences are indicated (Welch’s *t*-test; \**p* < 0.05, \*\**p* < 0.01).

**Table 1.**
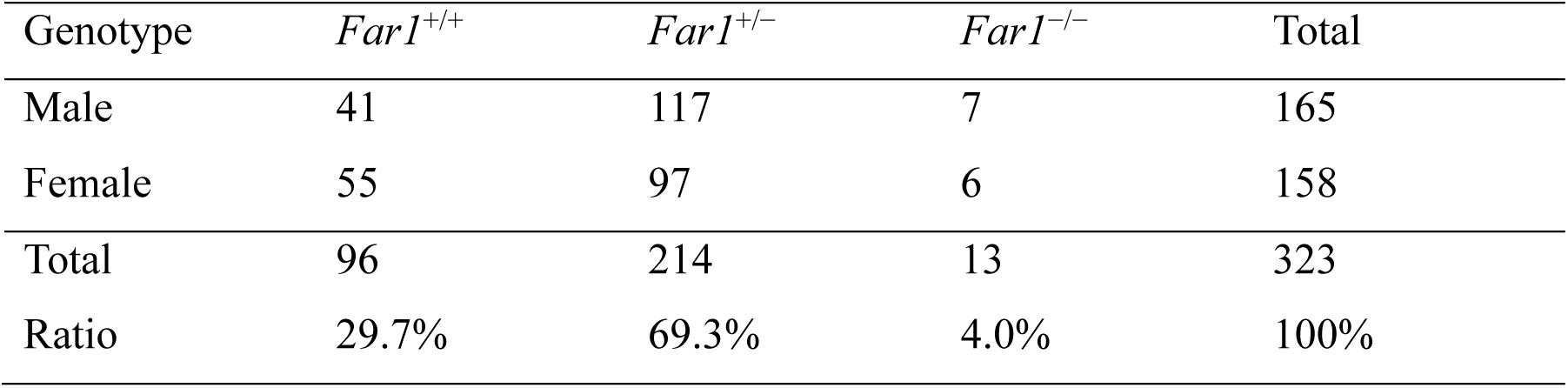
Number of *Far1*^+/+^, *Far1*^+/−^, and *Far1*^−/−^ mice at three weeks after birth.

**Table 2.**
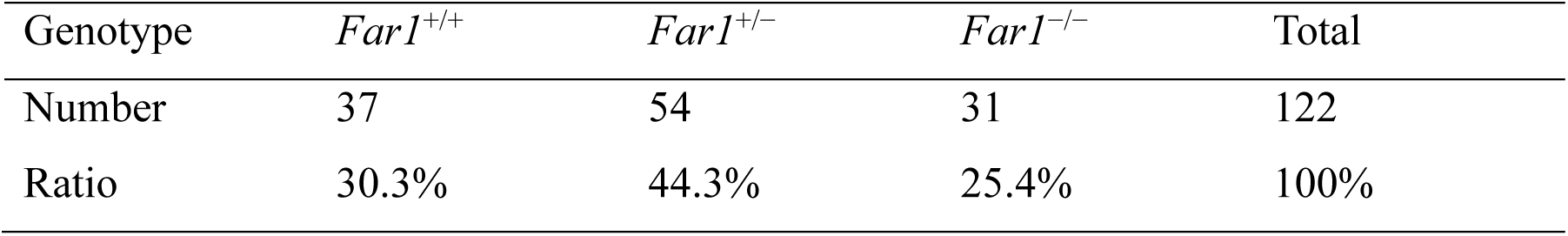
Number of *Far1*^+/+^, *Far1*^+/−^, and *Far1*^−/−^ mice at embryonic day 18.5.

Adult male *Far1* KO mice were normal in appearance (Fig. 4*C*). They had smaller body weights than WT mice, but this difference was not statistically significant (Fig. 4*C*). The testes of *Far1* KO mice were smaller than those of WT mice, with weights that were 40% of WT mouse testes (Fig. 4*D*). In WT mice, the epididymis appeared white because it contained mature sperms (22), but in *Far1* KO mice, it was rather translucent (Fig. 4*E*). The fertility status of male WT, heterozygous *Far1* KO, and *Far1* KO mice was examined by crossing them with female WT mice. Crossing with male WT or heterozygous *Far1* KO mice resulted in pregnancy and the delivery of offspring within one and a half months (Table 3). In contrast, crossing with male *Far1* KO mice did not result in pregnancy even after more than three months, and no offspring were produced. These results indicate that the male *Far1* KO mice are infertile.

**Table 3.**
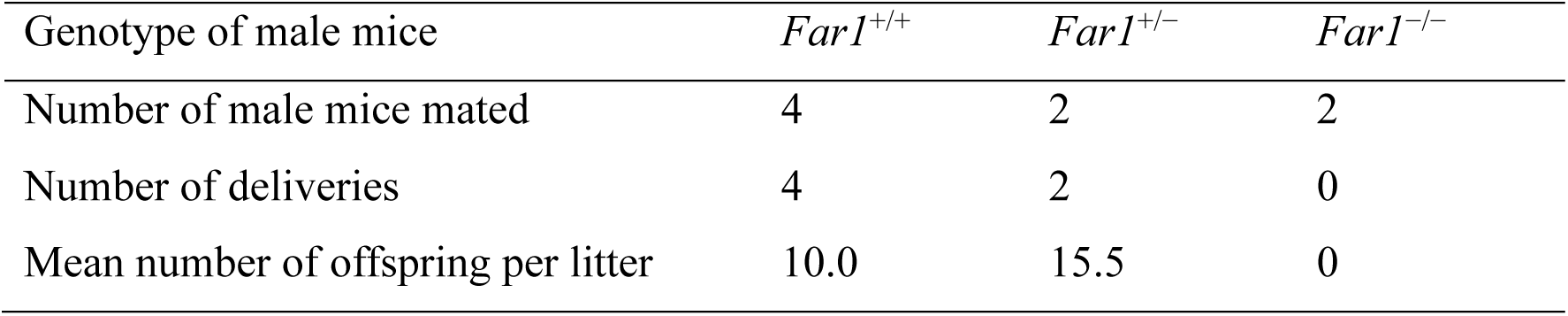
Number of offsprings obtained by mating male *Far1*^+/+^, *Far1*^+/−^, and *Far1*^−/−^ mice with WT female mice.

Histological analyses were performed on the epididymides and testes of WT and *Far1* KO mice by staining paraffin sections with hematoxylin-eosin. The epididymides of WT mice were full of sperms, whereas those of *Far1* KO mice contained none (Fig. 4*F*). In the seminiferous tubules in the testes of WT mice, spermatogonia, spermatocytes, and spermatids were readily observed, aligned from the basal lamina toward the center of the seminiferous tubules (Fig. 4*G*, black dashed ovals). In contrast, in *Far1* KO mice the cell positions were disorganized, and multinucleated cells were present in the seminiferous tubules (Fig. 4*G*, white arrowheads), suggesting impaired cytokinesis in the spermatogenic cells. In addition, there were vacuoles in the seminiferous tubules (Fig. 4*G*, asterisks). Sertoli cells were present only near or attached to the basal lamina of the seminiferous tubules in WT mice, while in *Far1* KO mice they were also ectopically present in the center of the tubule (Fig. 4*G*, black arrowheads).

The mRNA levels of pan-spermatogenic cell marker *Ddx4* (*DEAD box polypeptide 4*) (23) and Sertoli cell marker *Wt1* (*Wilms tumor 1*) (24) in the testes were examined via quantitative RT-PCR. The levels of *Ddx4* were reduced in *Far1* KO mice relative to WT mice, while the *Wt1* levels were comparable between the two groups (Fig. 4*H*). Next, to examine the differentiation of spermatogenic cells, we also quantified the mRNA levels of spermatogonium cell markers *Ccna1* (*Cyclin A1*) (25) and *Hsp70-2* (*heat shock protein 70*) (26) and spermatocyte markers *Prm1* (*Protamine 1*) (27) and *Hspa1l* (*Heat shock protein family A member 1 like*) (28). In *Far1* KO mice, the mRNA levels of *Ccna1* were significantly reduced to 43% of those in WT mice (Fig. 4*I*). The mRNA levels of *Hsp70-2* showed a similar reduction, although that decrease was not statistically significant. The mRNA levels of *Prm1* and *Hspa1l* in *Far1* KO mice were reduced to less than 5% of those in WT mice. Since the *Ccna1* and *Hsp70-2* genes are highly expressed in spermatogonia during the prophase of meiosis I, in which they are important (26, 29), these results indicate that, in *Far1* KO mice, the spermatocytes fail to complete meiosis I, resulting in the impaired differentiation of spermatocytes into spermatids. Combined, these results demonstrate that male *Far1* KO mice are infertile due to impaired spermatogenesis in the testes.

### Impaired production of seminolipids in *Far1* KO mice

To investigate the contribution of FAR1 to seminolipid synthesis, we extracted lipids from the testes of WT and *Far1* KO mice and analyzed the seminolipids using LC-MS/MS. In the *Far1* KO mice, seminolipids were essentially absent (Fig. 5*A*). Lipids were then separated via TLC, and glycolipids were detected using orcinol sulfate staining. The seminolipid band detected in WT mice was not entirely absent in the *Far1* KO mice but remained faintly detectable (Fig. 5*B*). This apparent discrepancy between the LC-MS/MS and TLC analyses (Fig. 5, *A* and *B*) may be explained by the presence of other lipids that were structurally similar to seminolipids in *Far1* KO mice. We hypothesized that these lipids may have been sulfogalactosyl-diacylglycerols (SGalDAGs), which have a 1-acyl moiety instead of a 1-alkyl moiety as in seminolipids (Fig. 5*C*). To test this hypothesis, we scraped the band from the TLC plate, extracted the lipids, and subjected them to LC-MS/MS analysis. Based on the structural analogy of seminolipids, we applied the following settings for the LC-MS/MS analysis: one acyl moiety was set as C16:0 while the other was allowed to vary, with chain lengths of C14– 26 and 0–6 double bonds (Fig. 5*C*). This analysis revealed that SGalDAGs were present in trace quantities in the testes of WT mice but substantially more abundant in *Far1* KO mice (42-fold the quantities in WT mice; Fig. 5*D*). The most abundant species was C16:0-C16:0, which comprised 99% and 91% of total SGalDAGs in WT and *Far1* KO mice, respectively. These results indicate that seminolipids are almost absent in *Far1* KO mice, but there are a compensatory increase in SGalDAGs.

**Figure 5.**
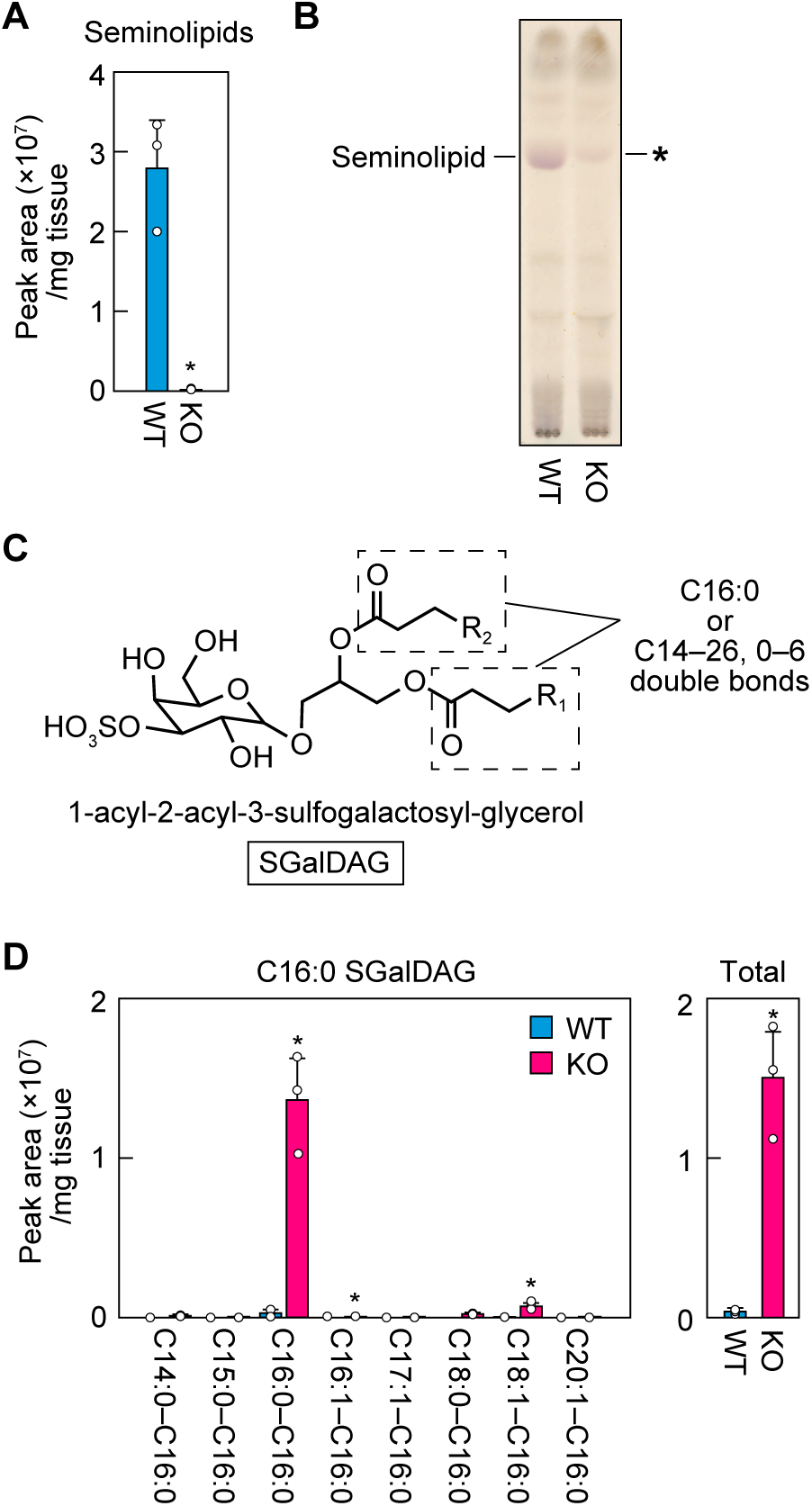
Large decrease in seminolipids and compensatory increase in SGalDAGs in *Far1* KO mice. *A*, lipids were extracted from the testes of 8-week-old male WT and *Far1* KO mice and seminolipids were quantified via LC-MS/MS. Values presented are means + SD (n = 3). Statistically significant differences are indicated (Welch’s *t*-test; \**p* < 0.05). *B*, lipids extracted from the testes of 8-week-old male WT and *Far1* KO mice were separated via TLC and glycolipids were detected via orcinol sulfate staining. *C*, structure of SGalDAG (1-acyl-2-acyl-3-sulfogalactosyl-glycerol). The ranges of chain lengths and degrees of unsaturation in 1-acyl and 2-acyl moieties (dotted rectangles), analyzed via LC-MS/MS, are indicated. *D*, lipids were extracted from the testes of 8-week-old male WT and *Far1* KO mice and SGalDAGs were quantified via LC-MS/MS. The quantity of each SGalDAG species and total quantity are presented. Values presented are means + SD (n = 3). Statistically significant differences are indicated (Welch’s *t*-test; \**p* < 0.05).

### Changes in the quantities of sulfatides, sphingomyelins, and ceramides in *Far1* KO mice

The enzymes that catalyze the last two steps in the synthesis of seminolipids, ceramide galactosyltransferase (CGT) and cerebroside sulfotransferase (CST), are also involved in the synthesis of sulfatides, the sulfated glycosphingolipids that are abundant in the brain, especially in myelin (Fig. 6*A*) (9, 30). Total lipids from the testis and brain of WT mice were separated via TLC and glycolipids were detected using orcinol sulfate staining. In the brain, non-hydroxy and 2-hydroxy forms of sulfatides and galactosylceramides were the predominant glycolipids (Fig. 6*B*). However, these glycolipids were not detected in the testes. To determine whether sulfatides were present in the testes, lipids extracted from the testes of WT and *Far1* KO mice were subjected to LC-MS/MS, which is more sensitive than TLC. This analysis revealed that trace quantities of sulfatide species with C16:0 as their acyl moiety were present (Fig. 6*C*). In *Far1* KO mice, the quantity of C16:0 sulfatide was 2.4 times higher than that in WT mice. The mRNA levels of *Cgt* and *Cst*, examined via quantitative RT-PCR, were not significantly different between WT and *Far1* KO mice (Fig. 6*D*) and thus were not correlated with the increases in the quantity of C16:0 sulfatide in *Far1* KO mice.

**Figure 6.**
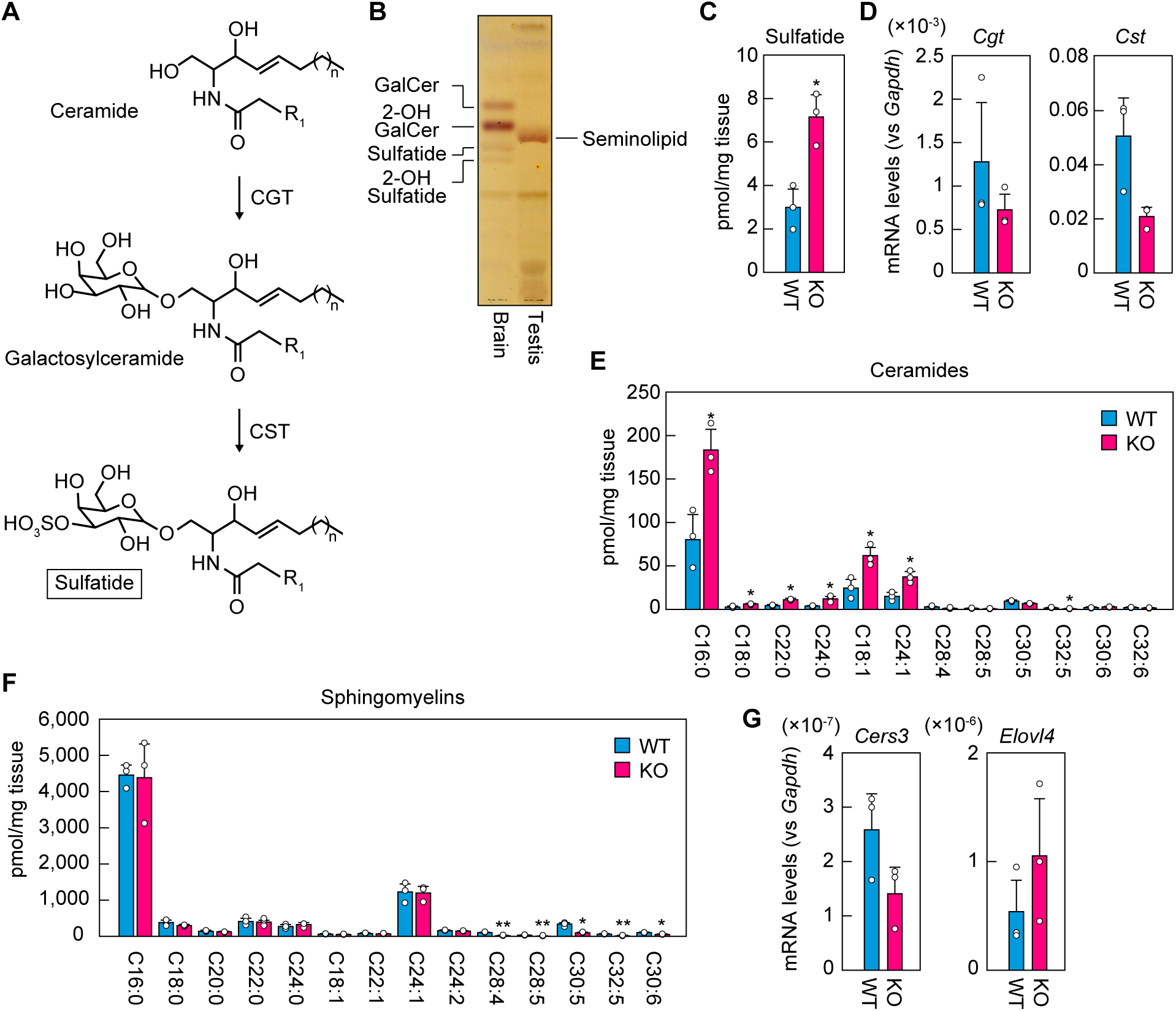
Changes in the quantities of sulfatides, sphingomyelins, and ceramides in the testes of *Far1* KO mice. *A*, synthetic pathway of sulfatide from ceramide. *B*, lipids were extracted from the brain (10-week-old, female) and testis (7-week-old, male) of C57BL/6 mice and separated via TLC. Glycolipids were detected via orcinol sulfate staining. GalCer, galactosylceramide; 2-OH, 2-hydroxy. *C*, lipids were extracted from the testes of 7-week-old male WT and *Far1* KO mice and sulfatides were quantified via LC-MS/MS. The quantities of sulfatide species containing a C16:0 acyl moiety are presented. Values presented are means + SD (n = 3). Statistically significant differences are indicated (Welch’s *t*-test; \**p* < 0.05). *D*, total RNAs were extracted from the testes of 8-week-old male WT and *Far1* KO mice and subjected to quantitative RT-PCR using specific primers for *Cgt*, *Cst*, or the housekeeping gene *Gapdh*. Values presented are means + SD of each mRNA quantity relative to *Gapdh* (n = 3). *E* and *F*, lipids were extracted from the testes of 8-week-old male WT and *Far1* KO mice and ceramides (*E*) and sphingomyelins (*F*) were quantified via LC-MS/MS. Values presented are means + SD of ceramide/sphingomyelin species containing the indicated acyl moiety (n = 3). Statistically significant differences are indicated (Welch’s *t*-test; \**p* < 0.05, \*\**p* < 0.01). *G*, total RNAs were extracted from the testes of 8-week-old male WT and *Far1* KO mice and subjected to quantitative RT-PCR using specific primers for *Elovl4*, *Cers3*, or the housekeeping gene *Gapdh*. Values presented are means + SD of mRNA quantities relative to *Gapdh* (n = 3).

In the testis, polyunsaturated fatty acids (PUFAs) such as C30:5 and C30:6 are found in the acyl moieties of sphingolipids such as ceramides and sphingomyelins (30). The ceramide synthase CERS3 is involved in the synthesis of these sphingolipids, and *Cers3* KO mice exhibit a deficiency in spermatogenesis (31). We measured ceramides and sphingomyelins in the testes of WT and *Far1* KO mice using LC-MS/MS. In the *Far1* KO mice, the levels of many ceramides and sphingomyelins containing ≥C28 PUFAs were significantly lower than in WT mice (Fig. 6, *E* and *F*). Conversely, the levels of ceramides containing C16–C24 saturated or monounsaturated fatty acids were higher. The mRNA levels of *Cers3* and *Elovl4*, the latter of which is involved in the synthesis of ≥C28 PUFAs, as examined using quantitative RT-PCR, were comparable between WT and *Far1* KO mice (Fig. 6*G*). Therefore, the low quantities of ceramides and sphingomyelins containing ≥C28 PUFAs in *Far1* KO mice may have reflected a deficiency in spermatogenesis.

## Discussion

It has not previously been determined which of the two mammalian FAR isozymes, FAR1 or FAR2, is involved in the synthesis of seminolipids in the testis. In this study, we observed that *Far1* was expressed in spermatogenic cells and *Far2* was not expressed in the testis (Fig. 2). *Far1* KO mice exhibited a loss of seminolipids in the testes, reduced numbers of spermatogenic cells and impaired spermatogenesis in the seminiferous tubules, reduced testicular weights, an absence of sperms in the epididymides, and male infertility (Figs. 4 and 5). *Far1* is therefore essential for seminolipid synthesis and spermatogenesis.

During our analyses of *Far1* KO mice, another group also generated *Far1* KO mice and reported the results of their analyses (19). Similar to the present study, they reported that these *Far1* KO mice exhibited partial postnatal lethality, reduced testicular weights, an absence of sperms, the presence of multinucleated spermatogenic cells, and reduced mRNA levels of genes expressed in spermatogonia and spermatocytes. However, while the authors of that study used LC-MS to show that plasmalogens (representatives of ether lipids) were almost absent, they did not analyze seminolipids. In this study, we found that seminolipids were absent in the testes of *Far1* KO mice (Fig. 5).

*Cgt* KO and *Cst* KO mice exhibited a loss of seminolipids, deficient spermatogenesis with multinucleated spermatogenic cells, and male infertility (8, 9); phenotypes similar to those in *Far1* KO mice. *Far1* is involved in the synthesis of ether lipids in general, including seminolipids and plasmalogens, whereas *Cgt* and *Cst* are involved in the synthesis of seminolipids but no other ether lipids; thus, seminolipids were identified as the ether lipids that are most important for spermatogenesis.

It has been established that *O*-C16:0/C16:0 is the most abundant seminolipid species in the testis (12–14). However, the diversity and detailed composition of the alkyl and acyl moieties, including chain lengths and degrees of unsaturation, in seminolipids have remained unknown. In this study, through comprehensive analyses using LC-MS/MS, we revealed the precise alkyl and acyl chain composition of seminolipids (Fig. 3). In addition, we examined SGalDAGs, which increased in *Far1* KO mice to compensate for the loss of seminolipids and sulfatides in the testis, and found that the predominant alkyl and/or acyl moieties in these lipids were C16:0 (Figs. 3, 5, and 6). In contrast, in the brain, the most abundant acyl moieties in sulfatides are C24:1, C24:0, and C22:0 (32, 33). Therefore, there appeared to be a testis-specific mechanism that promotes the selective incorporation of a C16:0 fatty acid/fatty alcohol into seminolipids, SGalDAGs, and sulfatides. Considering the acyl chain composition of sulfatides in the brain, the substrate specificities of CGT and CST cannot be limited to substrates with C16:0 alkyl/acyl chains. A possible mechanism is that the substrate specificity of CGT or CST is altered in favor of substrates containing C16:0 alkyl/acyl chains due to the unique membrane environment of the testis, which is enriched in PUFA-containing sphingolipids.

We observed multinucleated spermatogenic cells in the testes of *Far1* KO mice (Fig. 4). Therefore, it is possible that seminolipids are important for the formation and maintenance of intercellular bridges between spermatogenic cells, and that their deficiency causes the arrest of cytokinesis. The *O*-C16:0/C16:0 seminolipid is widely distributed in the seminiferous tubules of WT mice, as demonstrated by imaging MS (13). Immunohistochemical analysis using an antibody that recognizes sulfated glycolipids, including seminolipids, stained intercellular bridges in the seminiferous tubules (34). These observations suggest that seminolipids are components of intercellular bridges. The uniformity of seminolipids, which are composed almost entirely of *O*-C16:0/C16:0 species, may be necessary for the formation and maintenance of intercellular bridges. The *Far1* KO mice had increased levels of C16:0/C16:0 SGalDAGs in the testes but nevertheless exhibited multinuclear spermatogenic cells (Fig. 4), indicating the importance of the *O*-C16:0 moiety in seminolipids.

Multinucleated spermatogenic cells in seminiferous tubules have also been observed in mice with knockout of the genes involved in the synthesis of glycosphingolipids containing ultra-long-chain fatty acids (≥C26). These genes include the ceramide synthase *Cers3* and the glucosylceramide synthase *Ugcg* (31). Seminolipids may therefore cooperate with glycosphingolipids containing ultra-long-chain fatty acids in the formation and maintenance of intercellular bridges.

Sertoli cells perform multiple functions, such as providing physical support for spermatogenic cells, maintaining spermatogonia and their differentiation into spermatids, and forming the blood–testis barrier (2). In the *Far1* KO mice, cell positions were disorganized, and some Sertoli cells were ectopically located in the seminiferous tubules (Fig. 4). In the brain, sulfatides are involved in the formation of an adhesion structure between myelin and axons called the paranodal junction (9, 35). Therefore, seminolipids in the spermatogenic cells may also participate in the adhesion to Sertoli cells.

In summary, we found that FAR1 is involved in seminolipid synthesis and spermatogenesis in the testis. Most of the *Far1* KO mice died within a few days after birth, probably due to impaired production of plasmalogens—ether lipids that are abundant in the brain. Mutations in *FAR1* cause the inherited disease CSPSD, which causes neurological symptoms (15–18). *Far1* KO mice may be useful not only for elucidating the mechanism of spermatogenesis but also for investigating the pathogenic mechanisms of these neurological symptoms.

## Experimental procedures

### Mice

*Far1* KO mice were generated using the CRISPR-Cas9 system as follows. The guide RNA was designed to target the 20 bases adjacent to the protospacer-adjacent-motif sequence in exon 3 of *Far1*. A pair of oligonucleotides (5′-CACCGTCTTCCCCTCGTAGTATTCT-3′ and 5′-AAACAGAATACTACGAGGGGAAGAC-3′) containing the targeted sequence was annealed and cloned into the *Bbs*I site of the CRISPR/Cas9 vector pX330-U6-Chimeric_BB-CBh-hSpCas9 (Addgene, Watertown, MA, USA). The *Far1*-targeting plasmid was injected into fertilized eggs of C57BL/6J mice, and the injected eggs were transferred to the uteri of female C57BL/6J mice. To determine the genotypes of the offspring, genomic DNA from their tails was prepared and subjected to PCR using a pair of primers (5′-GGGATCCGTGAGTGATTTGTCTGATATGATCC-3′ and 5′-AATGCTTCACAAAATCCACACAAGC-3′) to amplify the DNA fragments containing the target sequence. The amplified DNA fragments were analyzed via agarose gel electrophoresis and Sanger sequencing. A founder mouse with a 22 bp deletion in exon 3 of *Far1* was obtained and crossed with C57BL/6J mice to establish a heterozygous *Far1* KO mouse line. Homozygous *Far1* KO mice were obtained by mating male and female heterozygous *Far1* KO mice. All the mice were housed under specific pathogen-free conditions at a room temperature of 23 ± 1 °C and a humidity of 50 ± 10%, with a 12 h light and 12 h dark cycle and water and food (Rodent Diet CE-2; CLEA Japan, Tokyo, Japan) available *ad libitum*. All animal experiments were approved by the Institutional Animal Care and Use Committee of Hokkaido University.

### Quantitative RT-PCR

After the testes of 8-week-old male mice had been dissected, they were immediately immersed in RNA*later* Stabilization Solution (Thermo Fisher Scientific, Waltham, MA, USA) and stored for ≥ 24 h at 4 °C. Total RNA was isolated using TRIzol Reagent (Thermo Fisher Scientific) and converted to first-strand cDNA using the PrimeScript II 1st Strand cDNA Synthesis Kit (Takara Bio, Shiga, Japan), following the manufacturer’s instructions. Real-time quantitative PCR was performed using the first-strand cDNA, KOD SYBR qPCR Mix (Toyobo, Osaka, Japan), and gene-specific primer pairs (Table 4) on the CFX96 Touch Real-Time PCR Detection System (Bio-Rad Laboratories, Hercules, CA). The mRNA levels were normalized with respect to *Gapdh*.

**Table 4.**
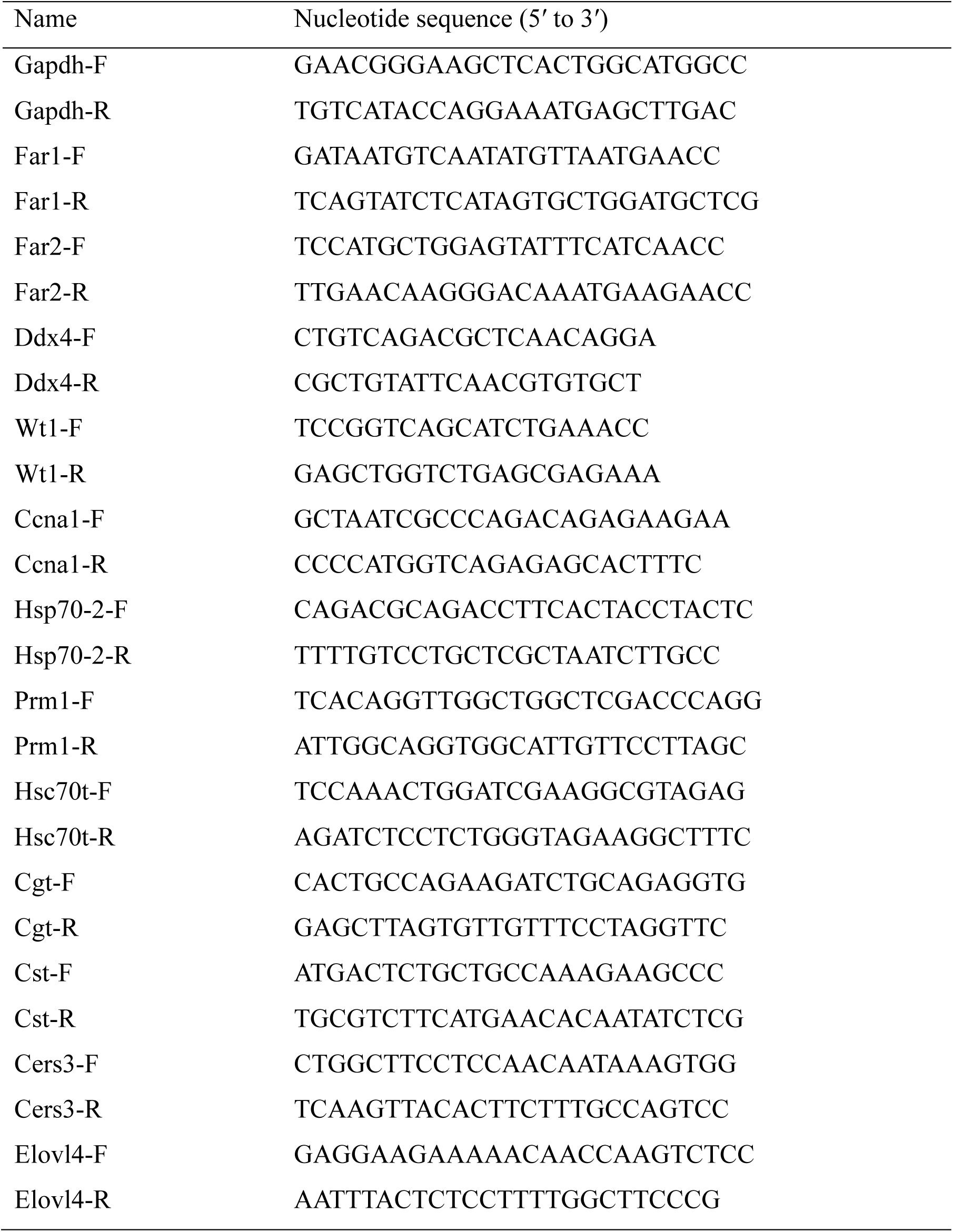
Oligonucleotide primers used in quantitative RT-PCR.

### *In situ* hybridization

To construct antisense RNA probes, *Far1* and *H2bc1* cDNA were amplified via PCR using the following primer pairs: for *Far1*, 5′-ATGGTTTCAATCCCAGAATACTACG-3′ and 5′-ACTACTTCATCAATATGCTTTCG-3′; and for *H2bc1*, 5′-ATGCCGGAGGTGGCGGTAAAGGGTG-3′ and 5′-TCACTTGGAGCTGGTGTACTTGGTG-3′. The amplified DNA fragments were cloned into the pGEM-T Easy Vector (Promega, Madison, WI, USA), and digoxigenin-labeled RNA probes were synthesized using the DIG RNA labeling mix (Merck, Darmstadt, Germany) and SP6 RNA polymerase (Merck).

*In situ* hybridization was performed as follows. Testes isolated from male mice at three days, two weeks, and six months after birth were frozen in Tissue-Tek OCT compound (Sakura Finetek Japan, Tokyo, Japan) at −80 °C. Samples were cut into 20 µm sections using a cryostat (CM3050S; Leica Biosystems, Wetzlar, Germany), attached to glass slides, fixed with 10% formaldehyde in PBS, and then hybridized with a digoxigenin-labeled *Far1* or *H2bc1* RNA probe. After washing, the hybridized probe was detected using alkaline phosphatase-conjugated anti-digoxigenin Fab fragments (Merck), followed by signal development for 6–24 h in a solution containing nitroblue tetrazolium and 5-bromo-4-chloro-3-indolyl phosphate (Merck). The samples were covered with glass coverslips using CC/Mount (Merck). Images were captured using a DM5000B light microscope (Leica Biosystems) equipped with a DFC295 digital color camera (Leica Biosystems).

### Separation of spermatogenic cells by FACS

The testes dissected from 12-week-old male mice were cut into small pieces and incubated with 1 mg/ml collagenase type I (Fujifilm Wako Pure Chemical, Osaka, Japan) and 0.5 units of DNase type I (Nippon Gene, Tokyo, Japan) in Dulbecco’s modified Eagle medium for 20 min at 33 °C with gentle rotation to digest extracellular matrices and DNA released from dead cells. The samples were then incubated with 0.25% trypsin (Fujifilm Wako Pure Chemical) for 20 min at 33 °C to disperse cells, and the reaction was terminated by adding 10% fetal bovine serum. After filtration through a 70 μm cell strainer, spermatogenic cells were stained with Hoechst 33342 solution (6 μg/million cells; Dojindo Laboratories, Kumamoto, Japan) for 30 min at 33 °C, followed by staining with 1 μg/ml propidium iodide solution (BioLegend, San Diego, CA, USA) for 10 min at room temperature. The samples were centrifuged (400*g*, 10 min, 4 °C), and the resulting pellets were suspended in PBS containing 2.5% fetal bovine serum, filtered through a 35 μm cell strainer, and sorted using the Cell Sorter SH800 (SONY, Tokyo, Japan). After excluding propidium-iodide–positive dead cells, spermatogenic cells were sorted into five major populations (spermatogonia, primary spermatocytes, secondary spermatocytes, round spermatids, and elongated spermatids) based on the intensity of two different wavelengths of fluorescence (Hoechst blue and Hoechst red) emitted by Hoechst33342 as described previously (36). Hoechst blue and Hoechst red were detected using 450/50 nm and 665/30 nm bandpass filters, respectively.

### Lipid extraction

Each testis dissected from the 8-week-old male mice was homogenized in 2.5 ml of chloroform/methanol/formic acid (100:200:1, v/v) containing 1 nmol of the C16:0 ceramide standard labeled with nine deuterium (*d*_9_), *N*-palmitoyl(*d*_9_) D-*erythro*-sphingosine (Avanti Research, Alabaster, AL, USA). The homogenate was centrifuged (1500*g*, 3 min, room temperature) and separated into supernatant and a pellet. The supernatant was recovered, and the pellet was subjected to a second extraction with 2.5 ml of chloroform/methanol/formic acid (100:200:1, v/v) containing 1 nmol of the *d*_9_*-*C16:0 ceramide standard and centrifuged as above. The supernatants from both extractions were combined. The combined sample was mixed with 3 ml of chloroform and 5.4 ml of water, vigorously mixed, and centrifuged (1500*g*, 3 min, room temperature). The resulting lower (organic) phase was recovered and dried.

For the alkaline hydrolysis of ester-linked lipids, the lipid sample, dissolved in 450 µl of chloroform/methanol (1:2, v/v), was mixed with 11.25 µl of 4 M potassium hydroxide in methanol and incubated for 1 h at 37 °C. The sample was neutralized by adding 11.25 µl of 4 M formic acid, followed by vigorous mixing. The sample was then mixed with 150 µl of chloroform and 270 µl of water, followed by centrifugation (20,600 *g*, 5 min, room temperature). The resulting organic phase was recovered and dried.

Brains dissected from 6–10-week-old female mice were chopped into small pieces. Ten milligrams of tissue was suspended in 450 µl of chloroform/methanol (1:2, v/v) in a tube containing zirconia beads (SARSTEDT, Nümbrecht, Germany) and vigorously mixed (4,500 rpm, 1 min, 4 °C, repeated twice) using a Micro Smash MS-100 (TOMY Seiko, Tokyo, Japan). The homogenate was centrifuged (20,600*g*, 5 min, room temperature) and separated into supernatant and a pellet. The supernatant was recovered and the pellet was subjected to a second extraction with 450 µl of chloroform/methanol (1:2, v/v) and centrifuged as above. The combined sample was mixed with 300 µl of chloroform and 540 µl of water, vigorously mixed, and centrifuged (20,600*g*, 5 min, room temperature). The resulting organic phase was recovered and dried.

Each population of spermatogenic cells, sorted as above, was centrifuged (400*g*, 10 min, 4 °C), and the pellet was suspended in 400 μl of chloroform/methanol (1:1, v/v) containing 10 pmol of the *d*_9_-C16:0 ceramide standard. After 180 μl of water had been added, the sample was vigorously mixed for 1 min and centrifuged (20,600*g*, 5 min, room temperature). The resulting organic phase was recovered and dried.

### LC-MS/MS analysis

For LC-MS/MS analysis, LC-coupled, triple quadrupole mass spectrometers (Xevo TQ-S and Xevo TQ-XS; Waters, Milford, MA, USA) were used. For the analyses of seminolipids, SGalDAGs, and sulfatides, the samples were dissolved in chloroform/methanol (1:2, v/v) and separated using a YMC-Triart C18 metal-free reversed-phase column (1.9 µm particle size, 2.1 mm inner diameter, 50 mm length; YMC, Kyoto Japan). The column temperature was set at 55 °C. The flow rate was set to 0.25 ml/min in a binary gradient system using mobile phase A (methanol/acetonitrile/water [1:1:3, v/v] containing 5 mM ammonium formate) and mobile phase B (2-propanol/water [49:1, v/v] containing 5 mM ammonium formate). The gradient steps were as follows: 0–1 min, 0% B; 1–5 min, linear gradient to 50% B; 5–24 min, linear gradient to 95% B; 25–25.1 min, step to 0% B; and 25.1–30 min, 0% B.

For the analysis of ceramides and sphingomyelins, the samples dissolved in chloroform/methanol (1:2, v/v) were separated using an ACQUITY UPLC CSH C18 reversed-phase column (1.7 µm particle size, 2.1 mm inner diameter, 100 mm length; Waters). The LC flow rate was 0.3 ml/min in a binary gradient system using mobile phase C (acetonitrile/water [3:2, v/v] containing 5 mM ammonium formate) and mobile phase D (acetonitrile/2-propanol [9:1, v/v] containing 5 mM ammonium formate). The gradient steps were as follows: 0 min, 40% D; 0–18 min, linear gradient to 100% D; 18–23 min, 100% D; 23–23.1 min, step to 40% D; and 23.1–25 min, 40% D.

Electrospray ionization was performed using the parameters listed in Table S1. MS/MS analysis was performed in multiple reaction monitoring mode using the *m*/*z* values of the precursor (Q1) and product (Q3) ions specific to each lipid species and optimized collision energies (Tables S2−S9). Data analysis was performed using the MassLynx software (Waters). The quantity of each seminolipid and SGalDAG species was presented as peak area because their standards were not commercially available. The quantity of each sulfatide, ceramide, and sphingomyelin species was calculated from its peak area as its ratio to the value of the corresponding C17:0 sulfatide (external standard; Avanti Research), *d*_9_-C16:0 ceramide (internal standard), and *d*_9_-C18:1 sphingomyelin (external standard; Avanti Research), respectively.

### Hematoxylin and eosin staining

The testes and epididymides of the 8-week-old WT and *Far1* KO mice were fixed with 3.7% formaldehyde in 0.1 M PBS (pH 7.4) for > 24 h at 4 °C. The fixed tissues were dehydrated, embedded in paraffin, cut into 4 µm-thick sections, deparaffinized, rehydrated, and stained with hematoxylin and eosin using an automated staining system (Tissue Tek DRS 2000; Sakura Finetek) as described previously (37). Bright-field images were captured using a Leica DM5000B microscope equipped with a DFC295 digital color camera (Leica Microsystems).

### Lipid analysis by TLC

Lipids (1 mg from the brain and 3–5 mg from the testis) were separated via TLC (Silica Gel 60 TLC plate, Merck) with methyl acetate/2-propanol/chloroform/methanol/0.25% calcium chloride in water (25:25:25:10:9, v/v) as the solvent system. The TLC plate was dried, sprayed with orcinol sulfate reagent (0.2% orcinol in 2.1 M aqueous sulfuric acid), dried again, and then heated to 100 °C to detect glycolipids.

## Data availability

All data generated or analyzed during this study are contained within the article.

## Supporting information

This article contains supporting information. Supplementary Tables S1–S9

## Author contributions

A. T., T. N., K. O., and T. T. investigation; T. S. writing–original draft; T. S. and A. K. supervision; A. K. writing–review and editing; A. K. conceptualization; A. K. project administration; T. S. and A. K. funding acquisition.

## Funding and additional information

This work was supported by KAKENHI (grant numbers: JP22H04986 to A.K. and JP23K24020 to T.S.) from the Japan Society for the Promotion of Science (JSPS) and by a grant from the Terumo Life Science Foundation (grant number; 24-Ⅲ4018).

## Abbreviations

CGT: ceramide galactosyltransferase
CST: cerebroside sulfotransferase
FAR: fatty acyl-CoA reductase
KO: knockout
MS/MS: tandem mass spectrometry
PUFA: polyunsaturated fatty acid
SGalDAG: sulfogalactosyl-diacylglycerol
WT: wild-type.

