## Supplemental information for "Fatty acyl-CoA reductase FAR1 is essential for testicular seminolipid synthesis, spermatogenesis, and male fertility"

**Table S1**

Parameters used in the LC-MS/MS analysis

| Lipids | Seminolipids and SGalDAGs | Sulfatides | Ceramides and Sphingomyelins |
|---|---|---|---|
| Ion mode | negative | negative | positive |
| Capillary voltage | 1.5 kV | 2.5 kV | 2.5 kV |
| Source temperature | 140 °C | 150 °C | 140 °C |
| Desolvation temperature | 650 °C | 250 °C | 650 °C |
| Cone gas flow | 150 L/h | 150 L/h | 150 L/h |
| Desolvation gas flow | 1200 L/h | 600 L/h | 1200 L/h |

**Table S2**

MS/MS settings for seminolipids containing a 1-alkyl C16:0 moiety using Xevo TQ-S

| 2-Acyl moiety | Precursor ion (Q1) $[M-H]^-$ | Product ion (Q3) | Cone voltage (V) | Collision energy (eV) |
|---|---|---|---|---|
| C14:0 | 767.5 | 539.3 | 40 | 60 |
| C15:0 | 781.6 | 539.3 | 40 | 60 |
| C16:0 | 795.6 | 539.3 | 40 | 60 |
| C17:0 | 809.6 | 539.3 | 40 | 60 |
| C18:0 | 823.7 | 539.3 | 40 | 60 |
| C19:0 | 837.7 | 539.3 | 40 | 60 |
| C20:0 | 851.7 | 539.3 | 40 | 60 |
| C21:0 | 865.7 | 539.3 | 40 | 60 |
| C22:0 | 879.8 | 539.3 | 40 | 60 |
| C23:0 | 893.8 | 539.3 | 40 | 60 |
| C24:0 | 907.8 | 539.3 | 40 | 60 |
| C25:0 | 921.8 | 539.3 | 40 | 60 |
| C26:0 | 935.9 | 539.3 | 40 | 60 |
| C14:1 | 765.5 | 539.3 | 40 | 60 |
| C15:1 | 779.6 | 539.3 | 40 | 60 |
| C16:1 | 793.6 | 539.3 | 40 | 60 |
| C17:1 | 807.6 | 539.3 | 40 | 60 |
| C18:1 | 821.6 | 539.3 | 40 | 60 |
| C19:1 | 835.7 | 539.3 | 40 | 60 |
| C20:1 | 849.7 | 539.3 | 40 | 60 |
| C21:1 | 863.7 | 539.3 | 40 | 60 |
| C22:1 | 877.7 | 539.3 | 40 | 60 |
| C23:1 | 891.8 | 539.3 | 40 | 60 |
| C24:1 | 905.8 | 539.3 | 40 | 60 |
| C25:1 | 919.8 | 539.3 | 40 | 60 |

| | | | | |
|---|---|---|---|---|
| C26:1 | 933.8 | 539.3 | 40 | 60 |

**Table S3**

MS/MS settings for seminolipids containing a 1-alkyl C16:0 moiety using Xevo TQ-XS

| 2-Acyl moiety | Precursor ion (Q1) $[M-H]^-$ | Product ion (Q3) | Cone voltage (V) | Collision energy (eV) |
|---|---|---|---|---|
| C14:0 | 767.6 | 539.3 | 90 | 60 |
| C15:0 | 781.7 | 539.3 | 90 | 60 |
| C16:0 | 795.7 | 539.3 | 90 | 60 |
| C17:0 | 809.7 | 539.3 | 90 | 60 |
| C18:0 | 823.8 | 539.3 | 90 | 60 |
| C19:0 | 837.8 | 539.3 | 90 | 60 |
| C20:0 | 851.8 | 539.3 | 90 | 60 |
| C21:0 | 865.8 | 539.3 | 90 | 60 |
| C22:0 | 879.9 | 539.3 | 90 | 60 |
| C23:0 | 893.9 | 539.3 | 90 | 60 |
| C24:0 | 907.9 | 539.3 | 90 | 60 |
| C25:0 | 921.9 | 539.3 | 90 | 60 |
| C26:0 | 936.0 | 539.3 | 90 | 60 |
| C14:1 | 765.6 | 539.3 | 90 | 60 |
| C15:1 | 779.7 | 539.3 | 90 | 60 |
| C16:1 | 793.7 | 539.3 | 90 | 60 |
| C17:1 | 807.7 | 539.3 | 90 | 60 |
| C18:1 | 821.7 | 539.3 | 90 | 60 |
| C19:1 | 835.8 | 539.3 | 90 | 60 |
| C20:1 | 849.8 | 539.3 | 90 | 60 |
| C21:1 | 863.8 | 539.3 | 90 | 60 |
| C22:1 | 877.8 | 539.3 | 90 | 60 |
| C23:1 | 891.9 | 539.3 | 90 | 60 |
| C24:1 | 905.9 | 539.3 | 90 | 60 |
| C25:1 | 919.9 | 539.3 | 90 | 60 |

| | | | | |
|---|---|---|---|---|
| C26:1 | 933.9 | 539.3 | 90 | 60 |
| C14:2 | 763.6 | 539.3 | 90 | 60 |
| C15:2 | 777.6 | 539.3 | 90 | 60 |
| C16:2 | 791.7 | 539.3 | 90 | 60 |
| C17:2 | 805.7 | 539.3 | 90 | 60 |
| C18:2 | 819.7 | 539.3 | 90 | 60 |
| C19:2 | 833.7 | 539.3 | 90 | 60 |
| C20:2 | 847.8 | 539.3 | 90 | 60 |
| C21:2 | 861.8 | 539.3 | 90 | 60 |
| C22:2 | 875.8 | 539.3 | 90 | 60 |
| C23:2 | 889.9 | 539.3 | 90 | 60 |
| C24:2 | 903.9 | 539.3 | 90 | 60 |
| C25:2 | 917.9 | 539.3 | 90 | 60 |
| C26:2 | 931.9 | 539.3 | 90 | 60 |
| C14:3 | 761.6 | 539.3 | 90 | 60 |
| C15:3 | 775.6 | 539.3 | 90 | 60 |
| C16:3 | 789.7 | 539.3 | 90 | 60 |
| C17:3 | 803.7 | 539.3 | 90 | 60 |
| C18:3 | 817.7 | 539.3 | 90 | 60 |
| C19:3 | 831.7 | 539.3 | 90 | 60 |
| C20:3 | 845.8 | 539.3 | 90 | 60 |
| C21:3 | 859.8 | 539.3 | 90 | 60 |
| C22:3 | 873.8 | 539.3 | 90 | 60 |
| C23:3 | 887.8 | 539.3 | 90 | 60 |
| C24:3 | 901.9 | 539.3 | 90 | 60 |
| C25:3 | 915.9 | 539.3 | 90 | 60 |
| C26:3 | 929.9 | 539.3 | 90 | 60 |
| C14:4 | 759.6 | 539.3 | 90 | 60 |
| C15:4 | 773.6 | 539.3 | 90 | 60 |

| | | | | |
|---|---|---|---|---|
| C16:4 | 787.6 | 539.3 | 90 | 60 |
| C17:4 | 801.7 | 539.3 | 90 | 60 |
| C18:4 | 815.7 | 539.3 | 90 | 60 |
| C19:4 | 829.7 | 539.3 | 90 | 60 |
| C20:4 | 843.7 | 539.3 | 90 | 60 |
| C21:4 | 857.8 | 539.3 | 90 | 60 |
| C22:4 | 871.8 | 539.3 | 90 | 60 |
| C23:4 | 885.8 | 539.3 | 90 | 60 |
| C24:4 | 899.8 | 539.3 | 90 | 60 |
| C25:4 | 913.9 | 539.3 | 90 | 60 |
| C26:4 | 927.9 | 539.3 | 90 | 60 |
| C14:5 | 757.6 | 539.3 | 90 | 60 |
| C15:5 | 771.6 | 539.3 | 90 | 60 |
| C16:5 | 785.6 | 539.3 | 90 | 60 |
| C17:5 | 799.6 | 539.3 | 90 | 60 |
| C18:5 | 813.7 | 539.3 | 90 | 60 |
| C19:5 | 827.7 | 539.3 | 90 | 60 |
| C20:5 | 841.7 | 539.3 | 90 | 60 |
| C21:5 | 855.8 | 539.3 | 90 | 60 |
| C22:5 | 869.8 | 539.3 | 90 | 60 |
| C23:5 | 883.8 | 539.3 | 90 | 60 |
| C24:5 | 897.8 | 539.3 | 90 | 60 |
| C25:5 | 911.9 | 539.3 | 90 | 60 |
| C26:5 | 925.9 | 539.3 | 90 | 60 |
| C14:6 | 755.6 | 539.3 | 90 | 60 |
| C15:6 | 769.6 | 539.3 | 90 | 60 |
| C16:6 | 783.6 | 539.3 | 90 | 60 |
| C17:6 | 797.6 | 539.3 | 90 | 60 |
| C18:6 | 811.7 | 539.3 | 90 | 60 |

| | | | | |
|---|---|---|---|---|
| C19:6 | 825.7 | 539.3 | 90 | 60 |
| C20:6 | 839.7 | 539.3 | 90 | 60 |
| C21:6 | 853.7 | 539.3 | 90 | 60 |
| C22:6 | 867.8 | 539.3 | 90 | 60 |
| C23:6 | 881.8 | 539.3 | 90 | 60 |
| C24:6 | 895.8 | 539.3 | 90 | 60 |
| C25:6 | 909.8 | 539.3 | 90 | 60 |
| C26:6 | 923.9 | 539.3 | 90 | 60 |

**Table S4**

MS/MS settings for seminolipids containing a 2-acyl C16:0 moiety using Xevo TQ-S

| 1-Alkyl moiety | Precursor ion (Q1) $[M-H]^-$ | Product ion (Q3) | Cone voltage (V) | Collision energy (eV) |
|---|---|---|---|---|
| C14:0 | 767.5 | 511.2 | 40 | 60 |
| C15:0 | 781.6 | 525.3 | 40 | 60 |
| C16:0 | 795.6 | 539.3 | 40 | 60 |
| C17:0 | 809.6 | 553.3 | 40 | 60 |
| C18:0 | 823.7 | 567.4 | 40 | 60 |
| C19:0 | 837.7 | 581.4 | 40 | 60 |
| C20:0 | 851.7 | 595.4 | 40 | 60 |
| C21:0 | 865.7 | 609.4 | 40 | 60 |
| C22:0 | 879.8 | 623.5 | 40 | 60 |
| C23:0 | 893.8 | 637.5 | 40 | 60 |
| C24:0 | 907.8 | 651.5 | 40 | 60 |
| C25:0 | 921.8 | 665.5 | 40 | 60 |
| C26:0 | 935.9 | 679.6 | 40 | 60 |
| C14:1 | 765.5 | 509.2 | 40 | 60 |
| C15:1 | 779.6 | 523.3 | 40 | 60 |
| C16:1 | 793.6 | 537.3 | 40 | 60 |
| C17:1 | 807.6 | 551.3 | 40 | 60 |
| C18:1 | 821.6 | 565.3 | 40 | 60 |
| C19:1 | 835.7 | 579.4 | 40 | 60 |
| C20:1 | 849.7 | 593.4 | 40 | 60 |
| C21:1 | 863.7 | 607.4 | 40 | 60 |
| C22:1 | 877.7 | 621.4 | 40 | 60 |
| C23:1 | 891.8 | 635.5 | 40 | 60 |
| C24:1 | 905.8 | 649.5 | 40 | 60 |
| C25:1 | 919.8 | 663.5 | 40 | 60 |

| | | | | |
|---|---|---|---|---|
| C26:1 | 933.8 | 677.5 | 40 | 60 |

**Table S5**

MS/MS settings for seminolipids containing a 2-acyl C16:0 moiety using Xevo TQ-XS

| 1-alkyl moiety | Precursor ion (Q1) $[M-H]^-$ | Product ion (Q3) | Cone voltage (V) | Collision energy (eV) |
|---|---|---|---|---|
| C14:0 | 767.6 | 511.2 | 90 | 60 |
| C15:0 | 781.7 | 525.3 | 90 | 60 |
| C16:0 | 795.7 | 539.3 | 90 | 60 |
| C17:0 | 809.7 | 553.3 | 90 | 60 |
| C18:0 | 823.8 | 567.4 | 90 | 60 |
| C19:0 | 837.8 | 581.4 | 90 | 60 |
| C20:0 | 851.8 | 595.4 | 90 | 60 |
| C21:0 | 865.8 | 609.4 | 90 | 60 |
| C22:0 | 879.9 | 623.5 | 90 | 60 |
| C23:0 | 893.9 | 637.5 | 90 | 60 |
| C24:0 | 907.9 | 651.5 | 90 | 60 |
| C25:0 | 921.9 | 665.5 | 90 | 60 |
| C26:0 | 936.0 | 679.6 | 90 | 60 |
| C14:1 | 765.6 | 509.2 | 90 | 60 |
| C15:1 | 779.7 | 523.3 | 90 | 60 |
| C16:1 | 793.7 | 537.3 | 90 | 60 |
| C17:1 | 807.7 | 551.3 | 90 | 60 |
| C18:1 | 821.7 | 565.3 | 90 | 60 |
| C19:1 | 835.8 | 579.4 | 90 | 60 |
| C20:1 | 849.8 | 593.4 | 90 | 60 |
| C21:1 | 863.8 | 607.4 | 90 | 60 |
| C22:1 | 877.8 | 621.4 | 90 | 60 |
| C23:1 | 891.9 | 635.5 | 90 | 60 |
| C24:1 | 905.9 | 649.5 | 90 | 60 |
| C25:1 | 919.9 | 663.5 | 90 | 60 |

| | | | | |
|---|---|---|---|---|
| C26:1 | 933.9 | 677.5 | 90 | 60 |
| C14:2 | 763.6 | 507.2 | 90 | 60 |
| C15:2 | 777.6 | 521.2 | 90 | 60 |
| C16:2 | 791.7 | 535.3 | 90 | 60 |
| C17:2 | 805.7 | 549.3 | 90 | 60 |
| C18:2 | 819.7 | 563.3 | 90 | 60 |
| C19:2 | 833.7 | 577.3 | 90 | 60 |
| C20:2 | 847.8 | 591.4 | 90 | 60 |
| C21:2 | 861.8 | 605.4 | 90 | 60 |
| C22:2 | 875.8 | 619.4 | 90 | 60 |
| C23:2 | 889.9 | 633.5 | 90 | 60 |
| C24:2 | 903.9 | 647.5 | 90 | 60 |
| C25:2 | 917.9 | 661.5 | 90 | 60 |
| C26:2 | 931.9 | 675.5 | 90 | 60 |
| C14:3 | 761.6 | 505.2 | 90 | 60 |
| C15:3 | 775.6 | 519.2 | 90 | 60 |
| C16:3 | 789.7 | 533.3 | 90 | 60 |
| C17:3 | 803.7 | 547.3 | 90 | 60 |
| C18:3 | 817.7 | 561.3 | 90 | 60 |
| C19:3 | 831.7 | 575.3 | 90 | 60 |
| C20:3 | 845.8 | 589.4 | 90 | 60 |
| C21:3 | 859.8 | 603.4 | 90 | 60 |
| C22:3 | 873.8 | 617.4 | 90 | 60 |
| C23:3 | 887.8 | 631.4 | 90 | 60 |
| C24:3 | 901.9 | 645.5 | 90 | 60 |
| C25:3 | 915.9 | 659.5 | 90 | 60 |
| C26:3 | 929.9 | 673.5 | 90 | 60 |
| C14:4 | 759.6 | 503.2 | 90 | 60 |
| C15:4 | 773.6 | 517.2 | 90 | 60 |

| | | | | |
|---|---|---|---|---|
| C16:4 | 787.6 | 531.2 | 90 | 60 |
| C17:4 | 801.7 | 545.3 | 90 | 60 |
| C18:4 | 815.7 | 559.3 | 90 | 60 |
| C19:4 | 829.7 | 573.3 | 90 | 60 |
| C20:4 | 843.7 | 587.3 | 90 | 60 |
| C21:4 | 857.8 | 601.4 | 90 | 60 |
| C22:4 | 871.8 | 615.4 | 90 | 60 |
| C23:4 | 885.8 | 629.4 | 90 | 60 |
| C24:4 | 899.8 | 643.4 | 90 | 60 |
| C25:4 | 913.9 | 657.5 | 90 | 60 |
| C26:4 | 927.9 | 671.5 | 90 | 60 |
| C14:5 | 757.6 | 501.2 | 90 | 60 |
| C15:5 | 771.6 | 515.2 | 90 | 60 |
| C16:5 | 785.6 | 529.2 | 90 | 60 |
| C17:5 | 799.6 | 543.2 | 90 | 60 |
| C18:5 | 813.7 | 557.3 | 90 | 60 |
| C19:5 | 827.7 | 571.3 | 90 | 60 |
| C20:5 | 841.7 | 585.3 | 90 | 60 |
| C21:5 | 855.8 | 599.4 | 90 | 60 |
| C22:5 | 869.8 | 613.4 | 90 | 60 |
| C23:5 | 883.8 | 627.4 | 90 | 60 |
| C24:5 | 897.8 | 641.4 | 90 | 60 |
| C25:5 | 911.9 | 655.5 | 90 | 60 |
| C26:5 | 925.9 | 669.5 | 90 | 60 |
| C14:6 | 755.6 | 499.2 | 90 | 60 |
| C15:6 | 769.6 | 513.2 | 90 | 60 |
| C16:6 | 783.6 | 527.2 | 90 | 60 |
| C17:6 | 797.6 | 541.2 | 90 | 60 |
| C18:6 | 811.7 | 555.3 | 90 | 60 |

| | | | | |
|---|---|---|---|---|
| C19:6 | 825.7 | 569.3 | 90 | 60 |
| C20:6 | 839.7 | 583.3 | 90 | 60 |
| C21:6 | 853.7 | 597.3 | 90 | 60 |
| C22:6 | 867.8 | 611.4 | 90 | 60 |
| C23:6 | 881.8 | 625.4 | 90 | 60 |
| C24:6 | 895.8 | 639.4 | 90 | 60 |
| C25:6 | 909.8 | 653.4 | 90 | 60 |
| C26:6 | 923.9 | 667.5 | 90 | 60 |

**Table S6**

MS/MS settings for SGalDAGs containing a acyl C16:0 moiety using Xevo TQ-S

| Acyl moiety | Precursor ion (Q1) $[M-H]^-$ | Product ion (Q3) | Cone voltage (V) | Collision energy (eV) |
|---|---|---|---|---|
| C14:0 | 781.6 | 553.4 | 40 | 60 |
| C15:0 | 795.7 | 553.4 | 40 | 60 |
| C16:0 | 809.7 | 553.4 | 40 | 60 |
| C17:0 | 823.7 | 553.4 | 40 | 60 |
| C18:0 | 837.8 | 553.4 | 40 | 60 |
| C19:0 | 851.8 | 553.4 | 40 | 60 |
| C20:0 | 865.8 | 553.4 | 40 | 60 |
| C21:0 | 879.8 | 553.4 | 40 | 60 |
| C22:0 | 893.9 | 553.4 | 40 | 60 |
| C23:0 | 907.9 | 553.4 | 40 | 60 |
| C24:0 | 921.9 | 553.4 | 40 | 60 |
| C25:0 | 935.9 | 553.4 | 40 | 60 |
| C26:0 | 950.0 | 553.4 | 40 | 60 |
| C14:1 | 779.6 | 553.4 | 40 | 60 |
| C15:1 | 793.7 | 553.4 | 40 | 60 |
| C16:1 | 807.7 | 553.4 | 40 | 60 |
| C17:1 | 821.7 | 553.4 | 40 | 60 |
| C18:1 | 835.7 | 553.4 | 40 | 60 |
| C19:1 | 849.8 | 553.4 | 40 | 60 |
| C20:1 | 863.8 | 553.4 | 40 | 60 |
| C21:1 | 877.8 | 553.4 | 40 | 60 |
| C22:1 | 891.8 | 553.4 | 40 | 60 |
| C23:1 | 905.9 | 553.4 | 40 | 60 |
| C24:1 | 919.9 | 553.4 | 40 | 60 |
| C25:1 | 933.9 | 553.4 | 40 | 60 |

| | | | | |
|---|---|---|---|---|
| C26:1 | 947.9 | 553.4 | 40 | 60 |
| C14:2 | 777.6 | 553.4 | 40 | 60 |
| C15:2 | 791.6 | 553.4 | 40 | 60 |
| C16:2 | 805.7 | 553.4 | 40 | 60 |
| C17:2 | 819.7 | 553.4 | 40 | 60 |
| C18:2 | 833.7 | 553.4 | 40 | 60 |
| C19:2 | 847.7 | 553.4 | 40 | 60 |
| C20:2 | 861.8 | 553.4 | 40 | 60 |
| C21:2 | 875.8 | 553.4 | 40 | 60 |
| C22:2 | 889.8 | 553.4 | 40 | 60 |
| C23:2 | 903.9 | 553.4 | 40 | 60 |
| C24:2 | 917.9 | 553.4 | 40 | 60 |
| C25:2 | 931.9 | 553.4 | 40 | 60 |
| C26:2 | 945.9 | 553.4 | 40 | 60 |
| C14:3 | 775.6 | 553.4 | 40 | 60 |
| C15:3 | 789.6 | 553.4 | 40 | 60 |
| C16:3 | 803.7 | 553.4 | 40 | 60 |
| C17:3 | 817.7 | 553.4 | 40 | 60 |
| C18:3 | 831.7 | 553.4 | 40 | 60 |
| C19:3 | 845.7 | 553.4 | 40 | 60 |
| C20:3 | 859.8 | 553.4 | 40 | 60 |
| C21:3 | 873.8 | 553.4 | 40 | 60 |
| C22:3 | 887.8 | 553.4 | 40 | 60 |
| C23:3 | 901.8 | 553.4 | 40 | 60 |
| C24:3 | 915.9 | 553.4 | 40 | 60 |
| C25:3 | 929.9 | 553.4 | 40 | 60 |
| C26:3 | 943.9 | 553.4 | 40 | 60 |
| C14:4 | 773.6 | 553.4 | 40 | 60 |
| C15:4 | 787.6 | 553.4 | 40 | 60 |

| | | | | |
|---|---|---|---|---|
| C16:4 | 801.6 | 553.4 | 40 | 60 |
| C17:4 | 815.7 | 553.4 | 40 | 60 |
| C18:4 | 829.7 | 553.4 | 40 | 60 |
| C19:4 | 843.7 | 553.4 | 40 | 60 |
| C20:4 | 857.7 | 553.4 | 40 | 60 |
| C21:4 | 871.8 | 553.4 | 40 | 60 |
| C22:4 | 885.8 | 553.4 | 40 | 60 |
| C23:4 | 899.8 | 553.4 | 40 | 60 |
| C24:4 | 913.8 | 553.4 | 40 | 60 |
| C25:4 | 927.9 | 553.4 | 40 | 60 |
| C26:4 | 941.9 | 553.4 | 40 | 60 |
| C14:5 | 771.6 | 553.4 | 40 | 60 |
| C15:5 | 785.6 | 553.4 | 40 | 60 |
| C16:5 | 799.6 | 553.4 | 40 | 60 |
| C17:5 | 813.6 | 553.4 | 40 | 60 |
| C18:5 | 827.7 | 553.4 | 40 | 60 |
| C19:5 | 841.7 | 553.4 | 40 | 60 |
| C20:5 | 855.7 | 553.4 | 40 | 60 |
| C21:5 | 869.8 | 553.4 | 40 | 60 |
| C22:5 | 883.8 | 553.4 | 40 | 60 |
| C23:5 | 897.8 | 553.4 | 40 | 60 |
| C24:5 | 911.8 | 553.4 | 40 | 60 |
| C25:5 | 925.9 | 553.4 | 40 | 60 |
| C26:5 | 939.9 | 553.4 | 40 | 60 |
| C14:6 | 769.6 | 553.4 | 40 | 60 |
| C15:6 | 783.6 | 553.4 | 40 | 60 |
| C16:6 | 797.6 | 553.4 | 40 | 60 |
| C17:6 | 811.6 | 553.4 | 40 | 60 |
| C18:6 | 825.7 | 553.4 | 40 | 60 |

| | | | | |
|---|---|---|---|---|
| C19:6 | 839.7 | 553.4 | 40 | 60 |
| C20:6 | 853.7 | 553.4 | 40 | 60 |
| C21:6 | 867.7 | 553.4 | 40 | 60 |
| C22:6 | 881.8 | 553.4 | 40 | 60 |
| C23:6 | 895.8 | 553.4 | 40 | 60 |
| C24:6 | 909.8 | 553.4 | 40 | 60 |
| C25:6 | 923.8 | 553.4 | 40 | 60 |
| C26:6 | 937.9 | 553.4 | 40 | 60 |

**Table S7**

MS/MS settings for sulfatides using Xevo TQ-XS

| Acyl moiety | Precursor ion (Q1) | | Product ion (Q3) | Cone voltage (V) | Collision energy (eV) |
|---|---|---|---|---|---|
| | $[M-H_2O-H]^-$ | $[M-H]^-$ | | | |
| C16:0 | 767.5 | 778.7 | 511.2 | 40 | 60 |
| C17:0 | 781.6 | 792.7 | 525.3 | 40 | 60 |
| C18:0 | 795.6 | 806.8 | 539.3 | 40 | 60 |
| C20:0 | 809.6 | 834.8 | 553.3 | 40 | 60 |
| C22:0 | 823.7 | 862.9 | 567.4 | 40 | 60 |
| C24:0 | 837.7 | 890.9 | 581.4 | 40 | 60 |
| C26:0 | 851.7 | 919.0 | 595.4 | 40 | 60 |
| C16:1 | 865.7 | 776.7 | 609.4 | 40 | 60 |
| C18:1 | 879.8 | 804.7 | 623.5 | 40 | 60 |
| C20:1 | 893.8 | 832.8 | 637.5 | 40 | 60 |
| C22:1 | 907.8 | 860.8 | 651.5 | 40 | 60 |
| C24:1 | 921.8 | 888.9 | 665.5 | 40 | 60 |
| C26:1 | 935.9 | 917.0 | 679.6 | 40 | 60 |
| 2-OH C16:0 | 765.5 | 794.7 | 509.2 | 40 | 60 |
| 2-OH C18:0 | 779.6 | 822.8 | 523.3 | 40 | 60 |
| 2-OH C20:0 | 793.6 | 850.8 | 537.3 | 40 | 60 |
| 2-OH C22:0 | 807.6 | 878.9 | 551.3 | 40 | 60 |
| 2-OH C24:0 | 821.6 | 906.9 | 565.3 | 40 | 60 |
| 2-OH C26:0 | 835.7 | 935.0 | 579.4 | 40 | 60 |
| 2-OH C16:1 | 849.7 | 792.7 | 593.4 | 40 | 60 |
| 2-OH C18:1 | 863.7 | 820.7 | 607.4 | 40 | 60 |
| 2-OH C20:1 | 877.7 | 848.8 | 621.4 | 40 | 60 |
| 2-OH C22:1 | 891.8 | 876.8 | 635.5 | 40 | 60 |
| 2-OH C24:1 | 905.8 | 904.9 | 649.5 | 40 | 60 |
| 2-OH C26:1 | 933.8 | 933.0 | 677.5 | 40 | 60 |

**Table S8**

MS/MS settings for ceramides using Xevo TQ-S

| Acyl moiety | Precursor ion (Q1) | | Product ion (Q3) | Cone voltage (V) | Collision energy (eV) |
|---|---|---|---|---|---|
| | $[M-H-H_2O]^-$ | $[M-H]^-$ | | | |
| C12:0 | 464.5 | 482.5 | 264.3 | 30 | 25 |
| C14:0 | 492.5 | 510.5 | 264.3 | 30 | 25 |
| C16:0 | 538.5 | 520.5 | 264.3 | 30 | 20 |
| C18:0 | 548.6 | 566.6 | 264.3 | 30 | 20 |
| C20:0 | 576.6 | 594.6 | 264.3 | 30 | 20 |
| C22:0 | 604.6 | 622.6 | 264.3 | 30 | 25 |
| C24:0 | 632.6 | 650.6 | 264.3 | 30 | 30 |
| C26:0 | 660.7 | 678.7 | 264.3 | 30 | 30 |
| C28:0 | 688.7 | 706.7 | 264.3 | 30 | 30 |
| C30:0 | 716.7 | 734.7 | 264.3 | 30 | 35 |
| C32:0 | 744.8 | 762.8 | 264.3 | 30 | 40 |
| C34:0 | 772.8 | 790.8 | 264.3 | 30 | 40 |
| C36:0 | 800.8 | 818.8 | 264.3 | 30 | 40 |
| C38:0 | 828.8 | 846.8 | 264.3 | 30 | 40 |
| C18:1 | 546.6 | 564.6 | 264.3 | 30 | 20 |
| C20:1 | 574.6 | 592.6 | 264.3 | 30 | 20 |
| C22:1 | 602.6 | 620.6 | 264.3 | 30 | 25 |
| C24:1 | 630.6 | 648.6 | 264.3 | 30 | 30 |
| C26:1 | 658.7 | 676.7 | 264.3 | 30 | 30 |
| C28:1 | 686.7 | 704.7 | 264.3 | 30 | 30 |
| C30:1 | 714.7 | 732.7 | 264.3 | 30 | 35 |
| C32:1 | 742.8 | 760.8 | 264.3 | 30 | 35 |
| C34:1 | 770.8 | 788.8 | 264.3 | 30 | 40 |
| C36:1 | 798.8 | 816.8 | 264.3 | 30 | 40 |
| C38:1 | 826.8 | 844.8 | 264.3 | 30 | 40 |

| | | | | | |
|---|---|---|---|---|---|
| C18:2 | 544.6 | 562.6 | 264.3 | 30 | 20 |
| C20:2 | 572.6 | 590.6 | 264.3 | 30 | 20 |
| C22:2 | 600.6 | 618.6 | 264.3 | 30 | 25 |
| C24:2 | 628.7 | 646.7 | 264.3 | 30 | 30 |
| C26:2 | 656.7 | 674.7 | 264.3 | 30 | 30 |
| C28:2 | 684.7 | 702.7 | 264.3 | 30 | 30 |
| C30:2 | 712.8 | 730.8 | 264.3 | 30 | 35 |
| C32:2 | 740.8 | 758.8 | 264.3 | 30 | 35 |
| C34:2 | 768.8 | 786.8 | 264.3 | 30 | 40 |
| C36:2 | 796.9 | 814.9 | 264.3 | 30 | 40 |
| C38:2 | 824.9 | 842.9 | 264.3 | 30 | 40 |
| C18:3 | 542.6 | 560.6 | 264.3 | 30 | 20 |
| C20:3 | 570.6 | 588.6 | 264.3 | 30 | 20 |
| C22:3 | 598.6 | 616.6 | 264.3 | 30 | 25 |
| C24:3 | 626.6 | 644.6 | 264.3 | 30 | 30 |
| C26:3 | 654.7 | 672.7 | 264.3 | 30 | 30 |
| C28:3 | 682.7 | 700.7 | 264.3 | 30 | 30 |
| C30:3 | 710.7 | 728.7 | 264.3 | 30 | 35 |
| C32:3 | 738.8 | 756.8 | 264.3 | 30 | 35 |
| C34:3 | 766.8 | 784.8 | 264.3 | 30 | 40 |
| C36:3 | 794.8 | 812.8 | 264.3 | 30 | 40 |
| C38:3 | 822.9 | 840.9 | 264.3 | 30 | 40 |
| C20:4 | 568.6 | 586.6 | 264.3 | 30 | 20 |
| C22:4 | 596.6 | 614.6 | 264.3 | 30 | 25 |
| C24:4 | 624.6 | 642.6 | 264.3 | 30 | 30 |
| C26:4 | 652.7 | 670.7 | 264.3 | 30 | 30 |
| C28:4 | 680.7 | 698.7 | 264.3 | 30 | 30 |
| C30:4 | 708.7 | 726.7 | 264.3 | 30 | 35 |
| C32:4 | 736.8 | 754.8 | 264.3 | 30 | 35 |

| | | | | | |
|---|---|---|---|---|---|
| C34:4 | 764.8 | 782.8 | 264.3 | 30 | 40 |
| C36:4 | 792.8 | 810.8 | 264.3 | 30 | 40 |
| C38:4 | 820.9 | 838.9 | 264.3 | 30 | 40 |
| C20:5 | 566.6 | 584.6 | 264.3 | 30 | 20 |
| C22:5 | 594.6 | 612.6 | 264.3 | 30 | 25 |
| C24:5 | 622.6 | 640.6 | 264.3 | 30 | 30 |
| C26:5 | 650.6 | 668.6 | 264.3 | 30 | 30 |
| C28:5 | 678.7 | 696.7 | 264.3 | 30 | 30 |
| C30:5 | 706.7 | 724.7 | 264.3 | 30 | 35 |
| C32:5 | 734.7 | 752.7 | 264.3 | 30 | 35 |
| C34:5 | 762.8 | 780.8 | 264.3 | 30 | 40 |
| C36:5 | 790.8 | 808.8 | 264.3 | 30 | 40 |
| C38:5 | 818.8 | 836.8 | 264.3 | 30 | 40 |
| C22:6 | 592.6 | 610.6 | 264.3 | 30 | 25 |
| C24:6 | 620.6 | 638.6 | 264.3 | 30 | 30 |
| C26:6 | 648.6 | 666.6 | 264.3 | 30 | 30 |
| C28:6 | 676.7 | 694.7 | 264.3 | 30 | 30 |
| C30:6 | 704.7 | 722.7 | 264.3 | 30 | 35 |
| C32:6 | 732.7 | 750.7 | 264.3 | 30 | 35 |
| C34:6 | 760.8 | 778.8 | 264.3 | 30 | 40 |
| C36:6 | 788.8 | 806.8 | 264.3 | 30 | 40 |
| C38:6 | 816.8 | 834.8 | 264.3 | 30 | 40 |

**Table S9**

MS/MS settings for sphingomyelins using Xevo TQ-S

| Acyl moiety | Precursor ion (Q1) $[M+H]^+$ | Product ion (Q3) | Cone voltage (V) | Collision energy (eV) |
|---|---|---|---|---|
| C12:0 | 647.5 | 184.1 | 50 | 60 |
| C14:0 | 675.6 | 184.1 | 50 | 60 |
| C16:0 | 703.6 | 184.1 | 50 | 60 |
| C18:0 | 731.7 | 184.1 | 50 | 60 |
| C20:0 | 759.8 | 184.1 | 50 | 60 |
| C22:0 | 787.8 | 184.1 | 50 | 60 |
| C24:0 | 815.9 | 184.1 | 50 | 60 |
| C26:0 | 843.9 | 184.1 | 50 | 60 |
| C28:0 | 872.0 | 184.1 | 50 | 60 |
| C30:0 | 900.0 | 184.1 | 50 | 60 |
| C32:0 | 928.1 | 184.1 | 50 | 60 |
| C34:0 | 956.1 | 184.1 | 50 | 60 |
| C36:0 | 984.2 | 184.1 | 50 | 60 |
| C38:0 | 1012.2 | 184.1 | 50 | 60 |
| C18:1 | 729.7 | 184.1 | 50 | 60 |
| C20:1 | 757.7 | 184.1 | 50 | 60 |
| C22:1 | 785.8 | 184.1 | 50 | 60 |
| C24:1 | 813.8 | 184.1 | 50 | 60 |
| C26:1 | 841.9 | 184.1 | 50 | 60 |
| C28:1 | 869.9 | 184.1 | 50 | 60 |
| C30:1 | 898.0 | 184.1 | 50 | 60 |
| C32:1 | 926.1 | 184.1 | 50 | 60 |
| C34:1 | 954.1 | 184.1 | 50 | 60 |
| C36:1 | 982.2 | 184.1 | 50 | 60 |
| C38:1 | 1010.2 | 184.1 | 50 | 60 |

| | | | | |
|---|---|---|---|---|
| C18:2 | 727.7 | 184.1 | 50 | 60 |
| C20:2 | 755.7 | 184.1 | 50 | 60 |
| C22:2 | 783.8 | 184.1 | 50 | 60 |
| C24:2 | 811.8 | 184.1 | 50 | 60 |
| C26:2 | 839.9 | 184.1 | 50 | 60 |
| C28:2 | 867.9 | 184.1 | 50 | 60 |
| C30:2 | 896.0 | 184.1 | 50 | 60 |
| C32:2 | 924.0 | 184.1 | 50 | 60 |
| C34:2 | 952.1 | 184.1 | 50 | 60 |
| C36:2 | 980.1 | 184.1 | 50 | 60 |
| C38:2 | 1008.2 | 184.1 | 50 | 60 |
| C18:3 | 725.7 | 184.1 | 50 | 60 |
| C20:3 | 753.7 | 184.1 | 50 | 60 |
| C22:3 | 781.8 | 184.1 | 50 | 60 |
| C24:3 | 809.8 | 184.1 | 50 | 60 |
| C26:3 | 837.9 | 184.1 | 50 | 60 |
| C28:3 | 865.9 | 184.1 | 50 | 60 |
| C30:3 | 894.0 | 184.1 | 50 | 60 |
| C32:3 | 922.0 | 184.1 | 50 | 60 |
| C34:3 | 950.1 | 184.1 | 50 | 60 |
| C36:3 | 978.1 | 184.1 | 50 | 60 |
| C38:3 | 1006.2 | 184.1 | 50 | 60 |
| C18:4 | 723.6 | 184.1 | 50 | 60 |
| C20:4 | 751.7 | 184.1 | 50 | 60 |
| C22:4 | 779.7 | 184.1 | 50 | 60 |
| C24:4 | 807.8 | 184.1 | 50 | 60 |
| C26:4 | 835.8 | 184.1 | 50 | 60 |
| C28:4 | 863.9 | 184.1 | 50 | 60 |
| C30:4 | 892.0 | 184.1 | 50 | 60 |

| | | | | |
|---|---|---|---|---|
| C32:4 | 920.0 | 184.1 | 50 | 60 |
| C34:4 | 948.1 | 184.1 | 50 | 60 |
| C36:4 | 976.1 | 184.1 | 50 | 60 |
| C38:4 | 1004.2 | 184.1 | 50 | 60 |
| C20:5 | 749.7 | 184.1 | 50 | 60 |
| C22:5 | 777.7 | 184.1 | 50 | 60 |
| C24:5 | 805.8 | 184.1 | 50 | 60 |
| C26:5 | 833.8 | 184.1 | 50 | 60 |
| C28:5 | 861.9 | 184.1 | 50 | 60 |
| C30:5 | 889.9 | 184.1 | 50 | 60 |
| C32:5 | 918.0 | 184.1 | 50 | 60 |
| C34:5 | 946.0 | 184.1 | 50 | 60 |
| C36:5 | 974.1 | 184.1 | 50 | 60 |
| C38:5 | 1002.2 | 184.1 | 50 | 60 |
| C22:6 | 775.7 | 184.1 | 50 | 60 |
| C24:6 | 803.8 | 184.1 | 50 | 60 |
| C26:6 | 831.8 | 184.1 | 50 | 60 |
| C28:6 | 859.9 | 184.1 | 50 | 60 |
| C30:6 | 887.9 | 184.1 | 50 | 60 |
| C32:6 | 916.0 | 184.1 | 50 | 60 |
| C34:6 | 944.0 | 184.1 | 50 | 60 |
| C36:6 | 972.1 | 184.1 | 50 | 60 |
| C38:6 | 1000.1 | 184.1 | 50 | 60 |
